# A miniaturized, high-throughput aqueous solvent-centric method for protein solubility screening

**DOI:** 10.1101/2025.06.16.659941

**Authors:** Adrian Svoboda, Marina Molineris, Theodora Tureckiova, Klára Hlouchová, Tomáš Pluskal, Teo Hebra

## Abstract

Efficient access to soluble recombinant proteins remains a major bottleneck in biochemical and structural studies. We describe an aqueous, solvent-centric, fully miniaturized 96-well workflow to screen extraction conditions that preserve soluble recombinant protein during lysis and clarification in a single working day. Liquid-nitrogen-frozen *E. coli* pellets are cryogenically bead-milled with stainless-steel beads, retaining the native intracellular milieu while ensuring uniform disruption. The resulting wet, frozen cell powder can therefore be extracted with user-defined solvent, enabling systematic exploration of pH, ionic strength, detergents, and chaotropes. Protein solubility is assessed by a 1 µL chromogenic anti-His dot-blot. We demonstrate the use of the protocol by solubilizing a set of highly challenging *de novo*-generated proteins and show that dot-blot intensity provides a practical semi-quantitative proxy for successful extraction of soluble proteins. We also provide experimentally supported guidelines on the influence of solvent reagents on subsequent steps of protein production, SDS-PAGE, and Ni-NTA purification. This workflow is compatible with upstream genetic solubility-enhancement, chassis- and cultivation-based strategies and enables direct transition from screening hits to scale-up. Because the workflow uses standard molecular biology equipment and inexpensive consumables, it can be readily adopted or automated in most laboratories.

## Introduction

Proper protein folding and solubility are prerequisites for most biochemical and structural investigations. Low protein solubility or aggregation may impair *in vitro* catalysis assays or subsequent steps for structural biology^1,2^. Although high-throughput *in vitro* enzymatic screening technologies have advanced rapidly,^3,4^ efficient production of soluble recombinant proteins has not kept pace.^5^ This remains a critical bottleneck in protein function discovery and structural characterization. Solubility depends on a broad spectrum of parameters (*e*.*g*. amino acid sequence,^6^ host genotype, expression cassette, culture medium, lysis conditions^7^). Although predictive methods are improving, their performance remains moderate^8,9^ and empirical solubility screening remains necessary. Therefore, for any given target or protein panel, the fastest route to soluble material is miniaturized, parallel expression screening^10,11^. Cultivation in 24- to 96-well plates enables systematic interrogation of the relevant variables impacting protein solubility.

Within the protein extraction workflow, cell disruption is critical: the transition from the intracellular milieu to an extraction solvent can alter protein solubility and promote aggregation^2^. Cell disruption is commonly achieved by chemical lysis, ultrasonication, or pressure-based methods such as French press or high-pressure homogenization. Pressure-based disruption is particularly effective for larger culture volumes, but both ultrasonication and pressure-based approaches are less convenient to standardize across many small samples in parallel and can cause sample warming during processing^12^. Chemical lysis is operationally simple, yet it often relies on additives (for example, detergents or strong denaturants) that can complicate downstream comparison of solubility across extraction conditions. Cryogenic bead milling disrupts liquid-nitrogen-frozen pellets with minimal warming, providing a gentle route to lysis that preserves the native cellular environment^13^. It is straightforward to parallelize in a 96-well format and can be scaled to match project scope^14^. Solvent composition is an equally powerful, yet underused, lever for improving solubilization. Solvent libraries are widely used for membrane-protein extraction^15^, immunoprecipitation^13^, and crystallization screens^16^, but they are rarely integrated into general recombinant protein production workflows^17^. We argue that extraction-solvent optimization should be implemented early in the screening procedure, and that it can be scaled to match any project scope or protein set.

Assessing soluble protein extraction constitutes a second bottleneck, as summarized by Baranowski *et al*.^5^ Among available readouts, anti-His dot-blot analysis combines “high parallelizability, low cost, and ease of sharing, and compatibility with downstream experiments; it is already widely used.^10,18,19^ Here we present a fully 96-well-plate compatible workflow that unites cryomilling, solvent-library screening, and anti-His solubility assay to assess the solubility space of recombinant proteins. We demonstrate the workflow on a set of cytosolic enzymes from three domains of life and five hard-to-express *de novo* proteins. The protocol leverages established upstream expression variables (host strain, induction temperature and timing, media composition, chaperone co-expression, and fusion tags) and enables medium- to high-throughput screening of extraction conditions to maximize soluble recovery. This provides a practical route to select conditions for subsequent functional and structural studies.

## Material and methods

### Growth Media, Strains and Plasmids

Chemicals used for media and extraction solvent preparation were purchased from Sigma-Aldrich, Duchefa Biochemie (Haarlem, Netherlands), Lach:ner (Neratovice, Czech Republic), or Penta Chemicals (Prague, Czech Republic). ZYM-5052 medium was prepared according to Studier et *al*.^20^ with ZY base: tryptone 1 % (v/w) yeast extract 0.5 % (v/w); M base (50 ×): 1.25 M Na_2_HPO_4_, 1.25 M KH_2_PO_4_, 2.5 M NH_4_Cl, 0.25 M Na_2_SO_4_; 5052 base (50 ×): glycerol 25 % (v/w), glucose 2.5 % (v/w), α-lactose 10 % (v/w); 1 M MgSO_4_; trace elements (1000 ×): 50 mM FeCl_3_; 20 mM CaCl_2_; 10 mM MnCl_2_; 10 mM ZnSO_4_; 2 mM CoCl_2_, 2 mM CuCl_2_; 2 mM NiCl_2_; 2 mM Na_2_MoO_4_; 2 mM Na_2_SeO_3_; 2 mM H_3_BO_3._

Plasmids encoding His8-Trx-mRuby2 and His8-MBP-mRuby2 were generated by Golden Gate assembly in 10 µL reactions containing T4 DNA ligase buffer (1 µL), T4 DNA ligase (0.5 µL; M0202L, New England Biolabs), BsaI-HFv2 (0.5 µL; R3733L, New England Biolabs), pYTK034 (mRuby2; 0.5 µL; 20 fmol), p3Xpress_Eco backbone (Trx or MBP tag; in-house), and nuclease-free water to 10 µL. Reactions were performed in a ProFlex 3 × 32-well PCR system thermocycler (Applied Biosystems) with 25 cycles of 37 °C for 5 min and 16 °C for 5 min, followed by 60 °C for 30 min and 80 °C for 10 min. DH10β electrocompetent *E. coli* cells were used for cloning, and transformants were selected on lysogeny broth agar containing kanamycin.

BL21(DE3) electrocompetent *E. coli* cells were used for expression of proteins. Transformed cells were selected on lysogeny broth (LB) with kanamycin.

### Protein extraction

A 1 mL *E. coli* BL21(DE3) culture was centrifuged for 10 min at 5,000 × g and 4 °C and the supernatant was removed. Cell pellets were flash-frozen by immersion in liquid nitrogen. One 5 mm stainless-steel bead was added, and frozen samples were mechanically lysed using QIAGEN TissueLyser II equipped with TissueLyser Adapter Set 2 x 24 (cat. no. 69982) or TissueLyser Adapter Set 2 x 96 (cat. no. 69984). Samples were milled at 25 Hz for 30 s, then incubated on ice for 1 min. Then 200 μL of extraction solvent was added to each sample and samples were resuspended by vortexing for 5 s, followed by three cycles of bath sonication (5 s) with cooling on ice (15 s) between cycles. Sonication was carried out in a Transsonic TS540 unit, 35 kHz, 50 watts per liter. Sample extraction after sonication was carried out after 5 min at 4 °C. To recover only soluble protein samples, the supernatant was collected after centrifugation for 20 min at 18,000 × g and 4 °C.

### Protein Purification

For small-scale cultivation (2 mL) Ni-NTA Agarose resin slurry (20 μL) of (QIAGEN) was washed three times by centrifugation 5 min, 700 *g*, 25 °C and resuspended in 30 μL of extraction solvent. 90 μL of clarified lysate was added to the resin and gently mixed for 7 min at 4 C. Samples were centrifuged for 5 min, 700 *× g*, 4 °C and supernatant was removed and pellet was resuspended in 300 μL of extraction solvent and centrifuged for 5 min, 700 *g*, 4 °C. Supernatant was removed and proteins were recovered from resin by addition of 20 μL of extraction solvent with 500 mM imidazole, gentle mixing for 3 min at 4 °C and centrifuged for 5 min, 700 *× g*, 4 °C.

For large-scale production of His8-MBP-mRuby2, 200 mL of *E. coli* BL21(DE3) culture expressing His8-MBP-mRuby2 was centrifuged at 5,000 *× g* for 30 min at 4 °C. The pellet was washed once with 1× PBS and centrifuged again at 5,000 *× g* for 30 min at 4 °C. The pellet was resuspended in 4 mL of 1 M ammonium acetate and transferred into two 10 mL TissueLyser II adapters for cryomilling. The adapters were flash-frozen in liquid nitrogen and the pellets were pulverized for 30 s at 25 Hz. The resulting powder was resuspended to a total volume of 40 mL with 1 M ammonium acetate (20 mL per adapter), briefly pulse-sonicated, and centrifuged at 15,000 *× g* for 30 min at 4 °C. The clarified lysate (in 1 M ammonium acetate) containing His8-MBP-mRuby2 was purified on a Ni-NTA column (4 mL resin). The protein was eluted in 25 mM Tris-HCl (pH 7.5) and 100 mM NaCl with stepwise imidazole concentrations of 50 mM, 250 mM, and 500 mM (12 mL each). Red fluorescent fractions were pooled and further purified on amylose resin (3 mL). Elution was performed in 25 mM Tris-HCl (pH 7.5) and 100 mM NaCl with stepwise maltose concentrations of 1, 2, 3, 5, and 10 mM. Fractions containing His8-MBP-mRuby2 were combined and concentrated using an Amicon centrifugal filter (10 kDa molecular weight cutoff; 4 mL device) to 0.5 mL (19 mg/mL). The sample was further purified by size-exclusion chromatography on a Superdex 200 Increase 10/300 GL column (Cytiva). Peak fractions corresponding to His8-MBP-mRuby2 were pooled and concentrated to 18 mg.mL^-1^ in the respective extraction solvent (see Figure 2). Protein concentrations were determined by absorbance at 280 nm.

**Figure 1.**
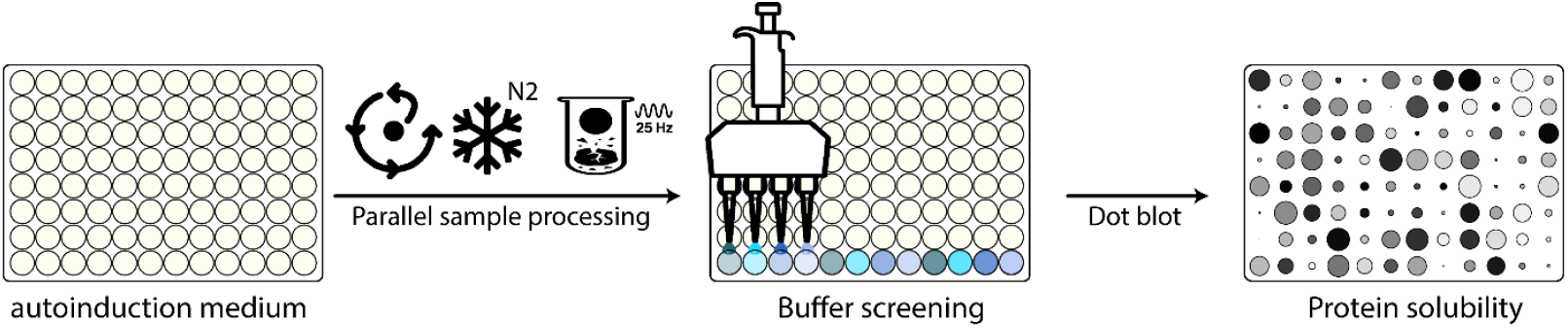
Workflow for quick assessment of protein solubility. Host cells for protein production are cultivated in 96-well plates overnight in auto-induction medium before being centrifuged, the supernatant removed, the pellets frozen in liquid nitrogen, and then mechanically lysed at 25 Hz with stainless-steel beads. Lysates are subsequently extracted with a set of solvents, and protein solubility is assessed by dot-blot.

**Figure 2.**
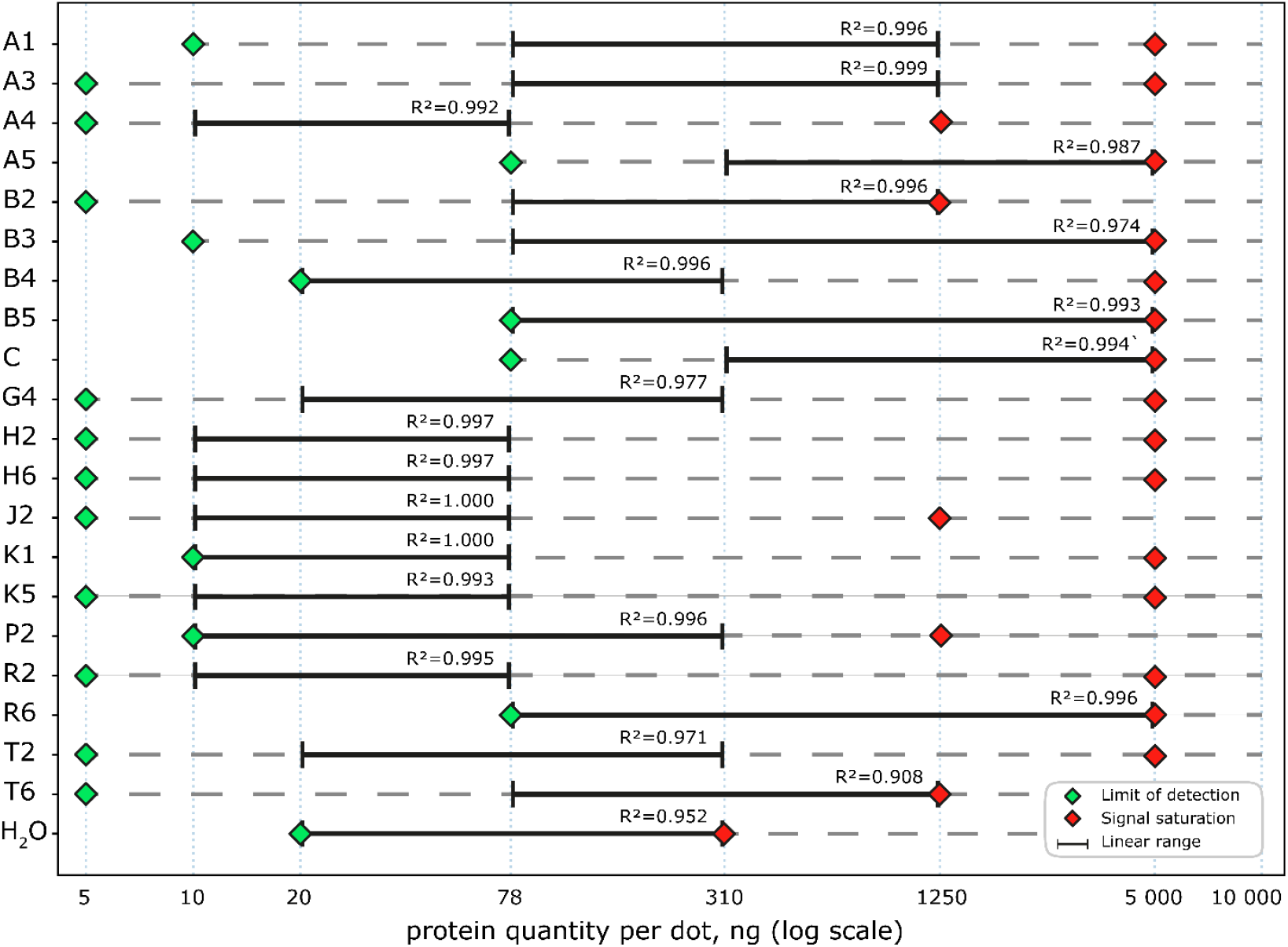
Semi-quantitative characterization of solvent effect on the dot-blot assay. For each solvent condition, the estimated linear dynamic range (black bars) was determined with corresponding R^2^. Green diamonds mark the limit of detection, and red diamonds indicate signal saturation. Protein quantities per dot are shown on a logarithmic scale.

### SDS-PAGE

For SDS-PAGE analysis, protein-containing solutions were denatured at 95 °C for 5 min and mixed with 2 × SDS loading buffer.

Samples were loaded in a 10 % acrylamide gel prepared with TGX FastCast acrylamide kit (Bio-Rad) and the electrophoresis was run for 40 min, 150 V. After electrophoresis, gel was washed with distilled water, covered with Coomassie Brilliant Blue, microwaved 20 sec, stained for 30 min at room temperature and was destained 2 h with MeOH/H_2_O/Acetic Acid (50:40:10) destaining solution at room temperature.

### Dot-blot

For dot-blot analysis, 1 μL of sample was transferred onto nitrocellulose membrane (10600012, cytiva) and dried for 30 min. The nitrocellulose membrane was blocked for 1 h in phosphate-buffered saline containing 5% milk, then incubated for 1 h with anti-polyhistidine–peroxidase antibody in the same buffer (A7058-1VL, Sigma Aldrich). The membrane was washed three times with 1 × PBS for 5 min and was visualized using 1 mL of 3,3′,5,5′-Tetramethylbenzidine (T0565-100mL, Sigma Aldrich) for 5 min.

### Fluorescence measurement

Fluorescence was measured using Spark^®^ Multimode Microplate Reader (Tecan) with the following parameters: excitation wavelength: 559 nm, excitation bandwidth: 15 nm, emission wavelength: 600 nm, emission bandwidth: 15 nm, Number of flashes: 30, Integration time: 40 µs, Gain optimal.

### Expression of terpene synthases

Active terpene synthases from previous work^21^ were cloned into p3Xpress_Eco (in-house) using Golden Gate assembly and transformed in *E. coli* BL21(DE3), as described in the Growth Media, Strains and Plasmids.

Three independent colonies were selected and inoculated in 2 mL of LB with kanamycin in a 24 deep-well plates (CR1426, Enzyscreen, Hamburg, Germany, Netherlands) sealed with AeraSeal™ (Excel Scientific, Victorville, California), at 37 °C, 800 rpm on an Eppendorf ThermoMixer® C (Eppendorf, Hamburg, Germany) for 3 h before induction with 0.5 mM IPTG (I1401, Duchefa Biochemie). Temperature was reduced to 18 °C and the induced cells were cultivated for 18 h.

### Amorphadiene synthase in vitro assay

Amorphadiene synthase (AMS) activity was assayed directly from clarified lysate obtained after cryomilling and extraction. Reactions were performed in a final volume of 250 µL by mixing 40 µL substrate solution (farnesyl pyrophosphate (FPP, Item No. 63250, Cayman Chemical), 0.2 mg·mL^−1^ in 25 mM NH_4_HCO_3_), 40 µL enzyme-containing clarified lysate (solvent B5), and 170 µL reaction buffer (50 mM HEPES (pH 7.0), 50 mM NaCl, 10 mM MgCl_2_, 100 mM urea, and 20% (v/v) glycerol). Reactions were incubated for 16 h at 30 °C with shaking at 1000 rpm. After incubation, products were extracted with ethyl acetate and the organic phase was analyzed by GC–MS using an Agilent 8890 GC system coupled to a LECO BTX MS. Data were processed in ChromaTOF (LECO), and product identity was assigned by spectral matching against the NIST MS database.

### Production of aggregation of prone proteins 1-5

For *de novo* generated proteins, an overnight *E. coli* BL21(DE3) culture expressing the target protein was used to inoculate 50 mL of lysogeny broth containing kanamycin and grown to an OD 600 nm of 0.5–0.6. Protein expression was induced with 0.5 mM IPTG and continued for 3 h at 30 °C. Cells were harvested by centrifugation at 10,000 *× g* for 5 min at 4 °C and divided into 2 mL aliquots to enable testing of multiple extraction conditions. Pellets were flash-frozen in liquid nitrogen.

### Fluorescence-lifetime imaging microscopy for aggregation prone proteins

To assess the localization and *in vivo* aggregation status, FLIM images of *de novo* generated proteins 1, 2, 5 were collected. The genes were subcloned to pETMF vector, providing expression with fusion to mTurquiose2 and mVenus^22^. Cultures of *E. coli* BL21(DE3) carrying the plasmids were induced at OD_600_ = 0.5 - 0.6 by IPTG at 0.5 mM concentration and incubated shaking for 18 hours at 25 °C. Cells washed with cold PBS were immobilized on poly-L-lysine coated glass cover slides and FLIM data were collected on Abberior Infinity STED microscope. Raw data was processed with in-house python script.^23^

### Fiji image processing

Dot intensities were quantified using Fiji (ImageJ)^24^. Images were cropped to the array area, the blue channel (mRuby2 experiments) or red channel (difficult-to-solubilize protein experiment) was extracted, converted to 8-bit grayscale, and inverted so that signal intensity increased with dot signal. Uneven background was corrected using the Subtract Background function with a rolling-ball radius of 100 pixels and the sliding paraboloid option. Dot intensities were measured using fixed-size circular regions of interest placed at predefined array positions. For each dot, integrated density was measured and background-corrected by subtracting the product of region area and mean background intensity. Within each membrane, values were normalized to the median signal of five internal control dots. To allow comparison across membranes, normalized values were scaled to the means of the control medians across all membranes.

### Data analysis

SDS-PAGE images were collected using an Azure 500 imager (near-infrared fluorescence with an orange tray). Dot-blot images were collected using an Azure 500 imager (true color). Images were processed in Fiji (cropping and contrast enhancement) using fiji software^24^. Raw data from fluorescence were processed using Rstudio and figures were generated using ggplot2 package^25^ and Adobe illustrator.

## Results

Our method is intended as a downstream, solvent-centric complement to upstream expression optimization. When soluble yield is limited by very low expression rather than post-lysis aggregation, upstream changes (for example lower induction temperature, altered induction duration, change in host strain) remain necessary. This solvent-centric workflow described here enables a post-cultivation, rapid, systematic screening of protein-solubility space and accommodates every common optimization parameter—host organism and strain, expression vector, fusion tag, growth medium, and chemical supplementation. Cells are cryomilled as nitrogen-frozen pellets, pulverized, extracted with user-defined solvent libraries, and evaluated for soluble expression by an anti-His dot-blot assay. For method development and solvent-compatibility benchmarking, we used mRuby2 as a robust soluble reporter protein; and its intrinsic red fluorescence allows rapid verification of target presence and normalization across workflow steps.

### Sample preparation workflow

We recommend 96-well deep-well plates for comprehensive solubility screening when many constructs or strains are tested in parallel against many solvent (Figure 1). For a single protein against multiple solvents, conventional baffled flasks suffice and 20 mL cultivation often allows screening up to 20 different solvents. Cultures are inoculated from fresh plates into ZYM-5052 auto-induction medium.^20^ After a 2 h incubation at 37 °C, the cultures are shifted to 18 °C for 14–18 h. Although auto-induction usually yields less protein than optimized IPTG induction, its gentle transcriptional activation and hand-off handling make it ideal for high-throughput campaigns. After protein production, cells are harvested at 5,000 × g for 10 min and the supernatant is discarded. Then, the pellets are plunged into liquid nitrogen. One stainless-steel bead is added per sample, and pellets are disrupted in a TissueLyser (30 s, 25 Hz). Following mechanical lysis, samples are warmed on ice for 1 min, extraction solvents are dispensed into each sample and suspensions are homogenized by three cycles of pulse sonication on a water-bath (5 s on / 15 s on ice). Lysates may be clarified by centrifugation immediately (30 min, 15,000 × g, 4 °C), but we found that a prolonged hold at 4 °C routinely increases the extraction yield.

### Dot-blot assay for rapid verification of protein solubility

Successful extraction of soluble target protein is assessed using a dot-blot with 1 µL of clarified lysate (or solvent control) per sample. Aliquots are spotted onto an 86 × 126 mm nitrocellulose membrane using the grid of a standard 96-well, 10 µL pipette-tip rack, which provides uniform spacing and reproducible spot geometry for up to 96 samples per membrane. After blotting, membranes are incubated with an anti-histidine antibody directly conjugated to horseradish peroxidase, and signal is developed using a chromogenic substrate. Under the conditions we used, spots become visible within 5 min after substrate addition and can be evaluated by eye without dedicated imaging equipment. It is worth noting that a broad range of anti-tag antibodies and conjugates is available, allowing the assay to be adapted to different laboratory setups and detection modalities. Overall, the procedure follows a conventional dot-blot workflow and provides a semi-quantitative readout of soluble expression within approximately 2 h.

### Solvent compatibility with downstream analyses

The extraction solvents used in this study were selected to span commonly used protein-extraction buffers (i.e. HEPES, Tris-HCl^26^ and Ammonium acetate^27,28^), modified with different concentrations of salt^29,30^ and additives (e.g. glycerol^31^, detergents^32^. DMSO^33^, ethanol^34^). They cover a representative range of conditions commonly used in biochemical assays but can be extended or modified as needed. Notably, the addition of detergent does not lead to any foaming during the extraction step. Because solubility screening is typically followed by protein characterization, we evaluated whether the extraction solvents interfered with three downstream readouts: (i) dot-blot detection that is our proposed assessment of extraction of soluble protein (Figure 2), (ii) SDS-PAGE gel electrophoresis followed by Coomassie Brilliant Blue staining (Figures S1–S9; Table 1) to assess that the right protein has been produced, and (iii) Ni-NTA affinity capture of histidine-tagged protein under native conditions to purify a protein of interest (Table 1).

**Table 1.** Effect of the extraction solvent composition on SDS-PAGE and Ni-NTA purification. SDS-PAGE: **+++**: sharp strong signal, +: peaks are spread or signal fainter, **-**: very faint signal of smear. W corresponds to the use of mQ water as a control. * yield is calculated based on the mRuby2 fluorescence of purified His8-MBP-mRuby2.

| Solvent Code | Solvent composition |  |  | SDS-PAGE | Purification yield, n = 2* |  |
| --- | --- | --- | --- | --- | --- | --- |
|  | Buffer | Salt | Additive |  |  |  |
| **A1** | Tris-HCl 50 mM; pH 8.5 | 50 mM NaCl | 10 % glycerol | +++ | **0.26** | **0.60** |
| **A3** | Tris-HCl 50 mM; pH 8.5 | 50 mM NaCl | 5 % ethanol | +++ | **0.41** | **0.44** |
| **A4** | Tris-HCl 50 mM; pH 8.5 | 50 mM NaCl | 1 % triton X100 | +++ | **0.58** | **0.67** |
| **A5** | Tris-HCl 50 mM; pH 8.5 | 50 mM NaCl | 100 mM Urea | +++ | **0.51** | **0.57** |
| **B2** | HEPES 50 mM; pH 7.0 | 50 mM NaCl | 5 % DMSO | +++ | **0.67** | **0.68** |
| **B3** | HEPES 50 mM; pH 7.0 | 50 mM NaCl | 5 % EtOH | +++ | **0.62** | **0.69** |
| **B4** | HEPES 50 mM; pH 7.0 | 50 mM NaCl | 1 % triton X100 | +++ | **0.76** | **0.68** |
| **B5** | HEPES 50 mM; pH 7.0 | 50 mM NaCl | 100 mM Urea | +++ | **0.81** | **0.64** |
| **C** | Tris-HCl 50 mM; pH 8.5 | 300 mM NaCl | 10 % glycerol | +++ | **0.45** | **0.66** |
| **G4** | Tris-HCl 20 mM, pH 8.0 | 400 mM NaCl | 1 % triton X100 | + | **0.71** | **0.76** |
| **H2** | Tris-HCl 20 mM, pH 8.0 | 0.4 M NH<sub>4</sub>Ac |  | + | **0.77** | **0.85** |
| **H6** | Tris-HCl 20 mM, pH 8.0 | 2.0 M NH<sub>4</sub>Ac |  | + | **0.74** | **0.82** |
| **J2** | Tris-HCl 20 mM, pH 8.0 | 0.4 M NH<sub>4</sub>Ac | 1 % triton X100 | + | **0.81** | **0.74** |
| **K1** |  | 2.0 M NH<sub>4</sub>Ac |  | - | **0.81** | **0.83** |
| **K5** |  | 1.5 M NH<sub>4</sub>Ac |  | +++ | **0.73** | **0.77** |
| **P2** | HEPES 50 mM; pH 7.4 | 200 mM NaCl | 1 % triton X100 | + | **0.91** | **0.83** |
| **R2** | HEPES 50 mM; pH 7.4 | 0.4 M NH<sub>4</sub>Ac |  | + | **0.77** | **0.85** |
| **R6** | HEPES 50 mM; pH 7.4 | 2.0 M NH<sub>4</sub>Ac |  | + | **0.61** | **0.80** |
| **T2** | HEPES 50 mM; pH 7.4 | 0.4 M NH<sub>4</sub>Ac | 1 % triton X100 | + | **0.87** | **0.81** |
| **T6** | HEPES 50 mM; pH 7.4 | 2.0 M NH<sub>4</sub>Ac | 1 % triton X100 | + | **0.76** | **0.79** |
| **W** | mQ water |  |  | +++ | **0.11** | **0.41** |

To assess solvent compatibility with the dot-blot readout independently of extraction efficiency, a single batch of His8-MBP-mRuby2 was produced and purified, then diluted into each test solvent at matched protein concentrations immediately before spotting as a serial dilution from 10,000 ng of protein per spot to 5 ng of protein per spot. To test whether solvent composition causes electrophoresis or staining artifacts (for example, altered migration, smearing, or reduced Coomassie staining), equal cell pellets were resuspended directly in the respective solvents and heat-denatured by boiling prior to addition of SDS loading buffer. This denaturing preparation serves as a stringent compatibility test and is not intended to quantify native soluble fractions. Samples were then analyzed by sodium dodecyl sulfate– polyacrylamide gel electrophoresis and visualized by standard Coomassie staining (Table 1, Figures S1– S9). In parallel, clarified lysates extracted using 20 solvents (prepared by cryomilling and cold extraction) were subjected to Ni-NTA purification under native conditions to assess recovery of histidine-tagged protein across solvent conditions (Table 1) and the yield of recovery was measured using mRuby2 fluorescence.

Most tested solvents supported detection of nanogram-scale quantities of protein and did not adversely affect the dot-blot readout relative to protein diluted in Milli-Q water. The main visible solvent-dependent effect was spot size, consistent with differences in surface tension. In addition, the tested solvents were broadly compatible with sodium dodecyl sulfate–polyacrylamide gel electrophoresis/Coomassie staining and Ni-NTA purification (Table 1). A concise summary of the few adverse interactions observed is provided in Table 1 to guide solvent choice and to indicate when a rapid solvent exchange is recommended before downstream analytical or preparative steps.

### Benchmarking the protocol on cytosolic enzymes from various domains

To test the protocol under realistic conditions, we selected a range of terpene synthases previously characterized by heterologous expression in yeast^21^. Those enzymes include bacterial and archaeal enzymes and one eukaryotic control (amorphadiene synthase, AMS, from flowering plants). Those enzymes displayed various behaviors on regular extraction methods (i.e. sonication) we attempted previously (from mg.mL^-1^ scale in soluble fraction to no protein in the soluble fraction). We subcloned the coding sequences into an in-house His-tag vector, inducible by IPTG and expressed the enzymes at 18 °C for 18 hours in 24 deep-well plate, 3 time 2 mL per construct. Then, we directly submitted the pellets to our cryomilling protocol using 3 different buffers and assessed the recovery of protein in the soluble fraction using dot blot (Figure 3). Finally, we verified that the enzyme retained activity after our extraction procedure by directly performing in vitro assay with the clarified lysate of buffer B5 containing amorphadiene synthase (an already characterized enzyme in prior literature^35^) and successfully detected its product by GC-MS.

**Figure 3.**
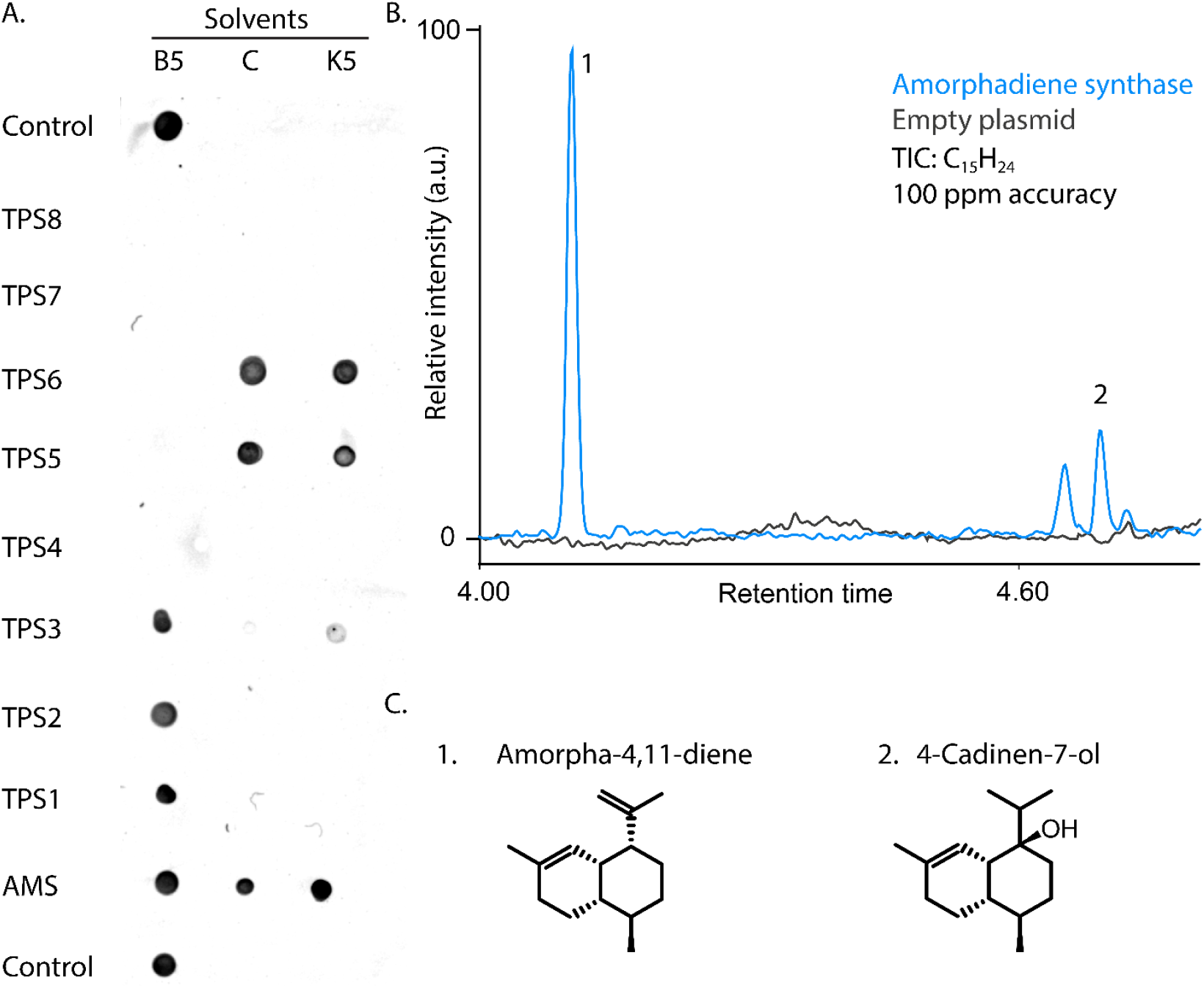
Functional validation of terpene synthase after extraction. (A) Anti-His dot blot analysis of soluble protein recovered after cryomilling and extraction in three solvent conditions (B5, C, K5). (B) GC–MS extracted ion chromatogram (TIC, C_15_H_24_) of ethyl acetate extracts of *in vitro* reactions using clarified lysate containing amorphadiene synthase (blue) or empty-vector control (black). (C) Structures of terpene products identified in the GC–MS analysis by NIST database search.

### Benchmarking the protocol on aggregation-prone proteins

After characterizing how extraction-solvent composition can affect downstream biochemical analyses and various cytosolic enzymes from eukaryotes, prokaryotes and archaea, we evaluated whether the protocol can rescue targets that are challenging to extract in the soluble fraction. We selected a set of *de novo* generated proteins that express mostly insolubly and respond poorly to standard expression-optimization strategies (Figure S10,11). In a previous attempt, these proteins were extracted from 500 mL cultures by chemical and enzymatic lysis (20 mL of buffer 1: 50 mM phosphate, pH 7.4; 150 mM sodium chloride; 2 mM magnesium chloride supplemented with lysozyme; Benzonase; and BugBuster), which consistently yielded insoluble material. For an initial demonstration of the present workflow, we screened five proteins (1-5) against five extraction solvents, alongside negative controls (*E. coli* induced cells carrying an empty vector and uninduced *E. coli* with pET30a(+) with protein 1) (Figure 4). Using our protocol, four of five proteins produced clear dot-blot signals in at least two extraction conditions. Notably, the extraction solvent used in the prior chemical-lysis workflow did not improve solubility in this assay, supporting the value of systematic solvent screening.

**Figure 4.**
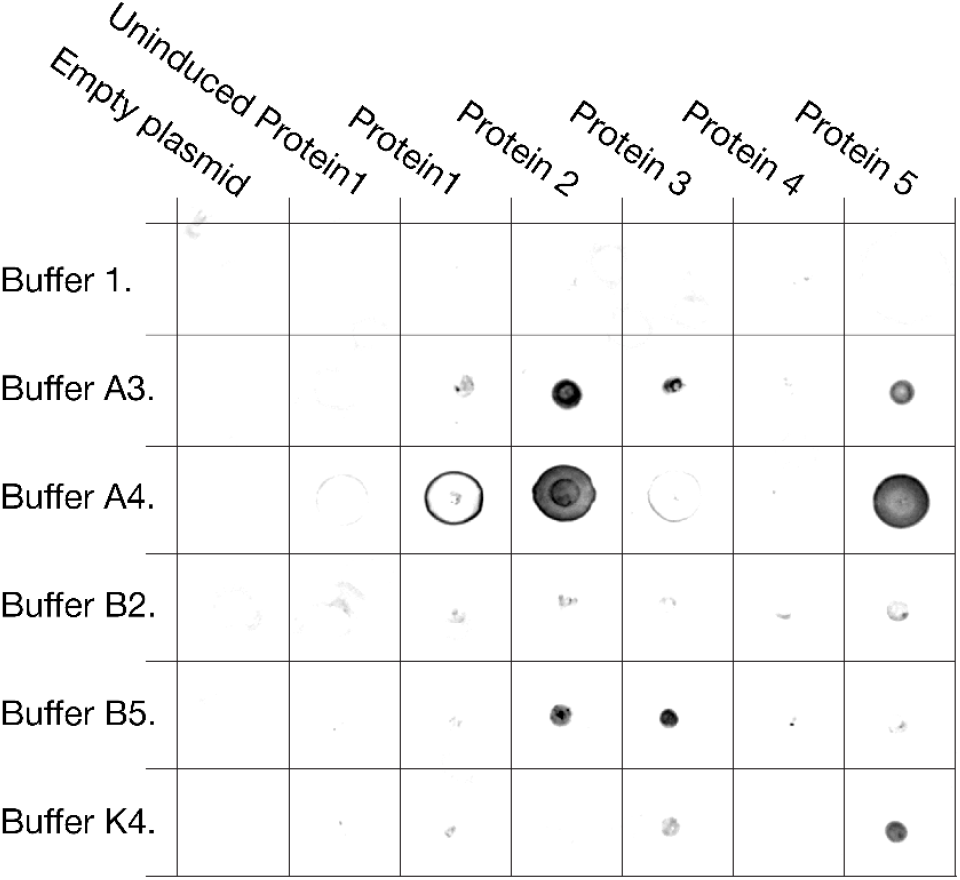
Dot-blot screening of difficult-to-solubilize proteins. Pellets from 2 mL *E. coli* cultures expressing proteins 1–5 were cryomilled and extracted with six solvents (Buffer 1, A3, A4, B2, B5, and K4). One dot corresponds to the clarified extract obtained with the indicated solvent. Negative controls were induced cells carrying an empty vector and uninduced cells carrying the plasmid encoding protein 1. Buffer compositions are listed in Table 1.

## Discussion

Our results show that a miniaturized workflow combining cultivation, cryomilling, solvent screening, and anti-His dot-blot enables parallel comparison of extraction conditions and supports rapid selection of conditions for scale-up. This integrated approach directly tackles the persistent bottleneck of obtaining soluble protein for downstream studies. Moreover, it integrates seamlessly with other parameters to increase protein solubility.^5,36^ A key innovation is the use of cryomilling for cell lysis. Cryogenic grinding of frozen cell pellets provided efficient and uniform disruption without the heat or shear stresses that can denature proteins during conventional lysis.^37^ This ensured that proteins were not prematurely aggregated by the lysis process. Moreover, it allows for high parallelization of the process.

The anti-His dot-blot is a rapid, convenient, and sensitive readout for extraction of soluble proteins. We reliably detected the target His-tagged protein directly from clarified lysates by spotting 1 μL samples onto a membrane, yielding a semi-quantitative “soluble/insoluble” signal from as little as 5-10 ng of protein load within approximately 2 h. This speed and throughput are difficult to achieve with SDS-PAGE and Western-blot workflows when screening dozens of conditions. The dot-blot signal provided an immediate readout for extraction of soluble proteins across solvents and helped guide final solvent selection for downstream experiments. Overall, the extraction solvents had only a modest impact on semi-quantitative dot-blot performance: limits of detection and linear dynamic ranges varied by no more than approximately 1.5 orders of magnitude across the tested conditions. The most pronounced solvent-dependent effect was surface tension, which altered spot spreading and therefore spot size. Accordingly, dot-blots can serve as a practical proxy for relative soluble protein extraction efficiency, but we still recommend confirming hit concentration using an independent concentration assay (for example, Bradford or absorbance at 280 nanometers) and a protein activity assay before selecting conditions for scale-up.

Not unexpectedly, the extraction solvent composition strongly affects both yield of extraction and post-lysis solubility—even for the highly soluble reporter mRuby2. This underscores the importance of the extraction solvent effect for proteins that are harder to express and extract at a yield high enough to proceed with further experiments such as crystallography. We found that certain additives had pronounced effects on SDS-PAGE and Ni-NTA purification. For example, solvents containing lower salt concentrations tend to provide a brighter signal in Coomassie blue mediated protein staining. However, for purification of the His-tagged protein, we observed the opposite effect: a higher salt concentration favors the binding of the proteins to the Ni-NTA resin, increasing the protein purification yield^37^. These observations highlight the importance of empirically screening a broad solvent space: each protein and subsequent experiment responds differently to solvent constituents, and there is no single “best” solvent for all targets and applications. Finally, we also demonstrated that our full procedure can be compatible with in vitro assay, directly from cell lysate, with enzymes retaining their enzymatic activity.

An additional strength of our strategy is its throughput and practicality for real-world protein expression optimization (Figure 3 and 4). The total sample requirement for each condition is small (1 mL of culture, 1 μL of extraction solvent for dot-blot), making the approach cost-effective. Importantly, we verified that the vast majority of the solvents in our library were compatible with standard downstream analyses. This means that hits from the screen can be translated to preparative scale with minimal adjustments. The present study validates the workflow on cytosolic, non-membrane recombinant proteins expressed in *E. coli*. We did not evaluate integral membrane proteins; although the workflow can, in principle, be adapted to user-defined targets, we do not draw conclusions about membrane-protein soluble recovery without dedicated validation.

By uniting upstream lysis through cryomilling with a downstream-friendly solvent screen, our workflow provides a streamlined path from expression to soluble protein ready for purification.

## Conclusion

We developed a high-throughput, small-volume method for screening recombinant protein solubility that integrates cryomilling-based cell lysis, a diverse set of extraction solvents, and a rapid dot-blot assay. This 96-well format workflow efficiently identifies optimal solvent conditions and maximizes the soluble yield of a target His-tagged protein. This method is effective as almost all tested solvent conditions are immediately compatible with standard purification and analysis steps such as Ni-NTA purification and SDS-PAGE analyses and *in vitro* assays, demonstrating a clear translation from screening leads to scalable protein production. The practical benefits (speed, low-cost, and the ability to pinpoint favorable conditions early) make this integrated method a valuable tool for any workflow aiming to produce challenging recombinant proteins.

## ASSOCIATED CONTENT

The following files are available free of charge. Supporting tables and figures (.pdf)

Raw data and raw imaged (.zip) The protocol

## AUTHOR INFORMATION

### Authors

**Adrian Svoboda - Institute of Organic Chemistry and Biochemistry of the Czech Academy of Sciences, 160 00 Prague, Czech Republic**

**Marina Molineris - Institute of Organic Chemistry and Biochemistry of the Czech Academy of Sciences, 160 00 Prague, Czech Republic**

**Theodora Tureckiova - Department of Cell Biology, Faculty of Science, Charles University, Prague 12800, Czech Republic;**

**Klára Hlouchová - Institute of Organic Chemistry and Biochemistry of the Czech Academy of Sciences, 160 00 Prague, Czech Republic; Department of Cell Biology, Faculty of Science, Charles University, Prague 12800, Czech Republic;**

**Tomáš Pluskal - Institute of Organic Chemistry and Biochemistry of the Czech Academy of Sciences, 160 00 Prague, Czech Republic;**

### Author Contributions

**A.S.: Investigation and validation. M.M.: Investigation. T.T.: Validation. K.H. Resources, funding. T.P.: Resources, funding, project administration, writing, and revisions. T.H.: Conceptualization, methodology, investigation, supervision, validation, writing, and revisions**.

### Funding Sources

**TP was supported by the Czech Science Foundation (GA CR) grant 21-11563M. TH was supported by the European Union’s Horizon Europe research and innovation program under the Marie Skłodowska-Curie grant agreement No. 101130799. Open access publishing facilitated by IOCB Prague as part of the ACS - CzechELib agreement**.

## ABBREVIATIONS

SDS-PAGE: sodium dodecyl sulfate–polyacrylamide gel electrophoresis
Ni-NTA: nickel Nitriloacetic acid
IPTG: Isopropyl β-D-1-thiogalactopyranoside
DMSO: Dimethyl sulfoxide
HEPES: 2-[4-(2-hydroxyethyl)piperazin-1-yl]ethanesulfonic acid
PBS: Phosphate-buffered saline

## AKNOWLEDGMENTS

We would like to thank Filip Buchel from Klára Hlouchová group for providing us the *de novo* generated protein 1-5.

We thank Dalibor Panek from Imaging Methods Core Facility at BIOCEV, institution supported by the MEYS CR (LM2023050 Czech-BioImaging), for their support with FLIM data acquisition and processing.

## Notes

### Competing Interest Statement

The authors have declared no competing interest.

### Summary of Updates

In the revised version, we strengthen our central claim by adding two new application datasets that go well beyond the original mRuby2 benchmarking: A panel of terpene synthases spanning all domains of life (bacteria, archaea, and eukaryotes), including functional validation for one model enzyme (new Figure 3). A set of de novo-designed, aggregation-prone proteins to test performance on difficult targets (new Figure 4). In addition to these new datasets demonstrating the protocol's value on realistic and challenging cases, we also tested and quantified how extraction-solvent composition influences the anti-histidine dot-blot readout, and we provide practical guidance for using the assay in a semi-quantitative manner (new Figure 2).

## REFERENCES

(01) Structural Genomics Consortium; China Structural Genomics Consortium; Northeast Structural Genomics Consortium; Gräslund, S.; Nordlund, P.; Weigelt, J.; Hallberg, B. M.; Bray, J.; Gileadi, O.; Knapp, S.; Oppermann, U.; Arrowsmith, C.; Hui, R.; Ming, J.; dhe-Paganon, S.; Park, H.-W.; Savchenko, A.; Yee, A.; Edwards, A.; Vincentelli, R.; Cambillau, C.; Kim, R.; Kim, S.-H.; Rao, Z.; Shi, Y.; Terwilliger, T. C.; Kim, C.-Y.; Hung, L.-W.; Waldo, G. S.; Peleg, Y.; Albeck, S.; Unger, T.; Dym, O.; Prilusky, J.; Sussman, J. L.; Stevens, R. C.; Lesley, S. A.; Wilson, I. A.; Joachimiak, A.; Collart, F.; Dementieva, I.; Donnelly, M. I.; Eschenfeldt, W. H.; Kim, Y.; Stols, L.; Wu, R.; Zhou, M.; Burley, S. K.; Emtage, J. S.; Sauder, J. M.; Thompson, D.; Bain, K.; Luz, J.; Gheyi, T.; Zhang, F.; Atwell, S.; Almo, S. C.; Bonanno, J. B.; Fiser, A.; Swaminathan, S.; Studier, F. W.; Chance, M. R.; Sali, A.; Acton, T. B.; Xiao, R.; Zhao, L.; Ma, L. C.; Hunt, J. F.; Tong, L.; Cunningham, K.; Inouye, M.; Anderson, S.; Janjua, H.; Shastry, R.; Ho, C. K.; Wang, D.; Wang, H.; Jiang, M.; Montelione, G. T.; Stuart, D. I.; Owens, R. J.; Daenke, S.; Schütz, A.; Heinemann, U.; Yokoyama, S.; Büssow, K.; Gunsalus, K. C. Protein Production and Purification. Nat Methods 2008, 5 (2), 135–146.

(02) Bondos, S. E.; Bicknell, A. Detection and Prevention of Protein Aggregation Before, During, and after Purification. Anal Biochem 2003, 316 (2), 223–231.

(03) Shepherd, R. A.; Fihn, C. A.; Tabag, A. J.; McKinnie, S. M. K.; Sanchez, L. M. “Need for Speed: High Throughput” - Mass Spectrometry Approaches for High-Throughput Directed Evolution Screening of Natural Product Enzymes. Nat Prod Rep 2025, 42 (6), 1037–1054.

(04) Markel, U.; Essani, K. D.; Besirlioglu, V.; Schiffels, J.; Streit, W. R.; Schwaneberg, U. Advances in Ultrahigh-Throughput Screening for Directed Enzyme Evolution. Chem Soc Rev 2020, 49 (1), 233–262.

(05) Baranowski, C.; Martin, H. G.; Oyarzún, D. A.; Spinner, A.; Desai, B.; Petzold, C. J.; Nikolados, E.-M.; Jaaks-Kraatz, S.; Gaber, A.; Chalkley, R. J.; Scannell, D.; Sevey, R.; Jewett, M. C.; Kelly, P. J.; DeBenedictis, E. A. Can Protein Expression Be “Solved”? Trends Biotechnol 2025, 43 (11), 2724–2742.

(06) Bhandari, B. K.; Gardner, P. P.; Lim, C. S. Solubility-Weighted Index: Fast and Accurate Prediction of Protein Solubility. Bioinformatics 2020, 36 (18), 4691–4698.

(07) Wingfield, P. T. Preparation of Soluble Proteins from Escherichia Coli. Curr Protoc Protein Sci 2014, 78, 6.2.1–6.2.22.

(08) Zhang, X.; Hu, X.; Zhang, T.; Yang, L.; Liu, C.; Xu, N.; Wang, H.; Sun, W. PLM_Sol: Predicting Protein Solubility by Benchmarking Multiple Protein Language Models with the Updated Escherichia Coli Protein Solubility Dataset. Brief Bioinform 2024, 25 (5), bbae404.

(09) Hon, J.; Marusiak, M.; Martinek, T.; Kunka, A.; Zendulka, J.; Bednar, D.; Damborsky, J. SoluProt: Prediction of Soluble Protein Expression in Escherichia Coli. Bioinformatics 2021, 37 (1), 23–28.

(10) Knaust, R. K.; Nordlund, P. Screening for Soluble Expression of Recombinant Proteins in a 96-Well Format. Anal Biochem 2001, 297 (1), 79–85.

(11) Berrow, N. S.; Büssow, K.; Coutard, B.; Diprose, J.; Ekberg, M.; Folkers, G. E.; Levy, N.; Lieu, V.; Owens, R. J.; Peleg, Y.; Pinaglia, C.; Quevillon-Cheruel, S.; Salim, L.; Scheich, C.; Vincentelli, R.; Busso, D. Recombinant Protein Expression and Solubility Screening in Escherichia Coli: A Comparative Study. Acta Crystallogr D Biol Crystallogr 2006, 62 (Pt 10), 1218–1226.

(12) Ferdous, S.; Dopp, J. L.; Reuel, N. F. Optimization of E. Coli Tip-Sonication for High-Yield Cell-Free Extract Using Finite Element Modeling. AIChE J 2021, 67 (10). 10.1002/aic.17389.

(13) Xie, S.; Saba, L.; Jiang, H.; Bringas, O. R.; Oghbaie, M.; Stefano, L. D.; Sherman, V.; LaCava, J. Multiparameter Screen Optimizes Immunoprecipitation. Biotechniques 2024, 76 (4), 145–152.

(14) Szmitkowska, A.; Pekárová, B.; Hejátko, J. A High-Throughput Strategy for Recombinant Protein Expression and Solubility Screen in Escherichia Coli : A Case of Sensor Histidine Kinase. Methods Mol Biol 2020, 2077, 19–36.

(15) Eshaghi, S.; Hedrén, M.; Nasser, M. I. A.; Hammarberg, T.; Thornell, A.; Nordlund, P. An Efficient Strategy for High-Throughput Expression Screening of Recombinant Integral Membrane Proteins. Protein Sci 2005, 14 (3), 676–683.

(16) Jancarik, J.; Kim, S.-H. Sparse Matrix Sampling: A Screening Method for Crystallization of Proteins. J. Appl. Crystallogr. 1991, 24 (4), 409–411. DOI: 10.1107/S0021889891004430.

(17) Peleg, Y.; Unger, T. Resolving Bottlenecks for Recombinant Protein Expression in E. Coli. Methods Mol Biol 2012, 800, 173–186.

(18) Vincentelli, R.; Canaan, S.; Offant, J.; Cambillau, C.; Bignon, C. Automated Expression and Solubility Screening of His-Tagged Proteins in 96-Well Format. Anal Biochem 2005, 346 (1), 77–84.

(19) Morris, M. A.; Bataglioli, R. A.; Mai, D. J.; Yang, Y. J.; Paloni, J. M.; Mills, C. E.; Schmitz, Z. D.; Ding, E. A.; Huske, A. C.; Olsen, B. D. Democratizing the Rapid Screening of Protein Expression for Materials Development. Mol. Syst. Des. Eng. 2023, 8 (2), 227–239.

(20) Studier, F. W. Protein Production by Auto-Induction in High Density Shaking Cultures. Protein Expr Purif 2005, 41 (1), 207–234.

(21) Samusevich, R.; Hebra, T.; Bushuiev, R.; Bushuiev, A.; Chatpatanasiri, R.; Kulhánek, J.; Čalounová, T.; Perković, M.; Engst, M.; Tajovská, A.; Sivic, J.; Pluskal, T. Discovery and Characterization of Terpene Synthases Powered by Machine Learning. bioRxiv, 2024. 10.1101/2024.01.29.577750.

(22) Aubel, M.; Buchel, F.; Heames, B.; Jones, A.; Honc, O.; Bornberg-Bauer, E.; Hlouchova, K. High-Throughput Selection of Human de Novo-Emerged sORFs with High Folding Potential. Genome Biol Evol 2024, 16 (4), evae069.

(23) https://github.com/IMCF-Biocev/Buchel_2025

(24) Schindelin, J.; Arganda-Carreras, I.; Frise, E.; Kaynig, V.; Longair, M.; Pietzsch, T.; Preibisch, S.; Rueden, C.; Saalfeld, S.; Schmid, B.; Tinevez, J.-Y.; White, D. J.; Hartenstein, V.; Eliceiri, K.; Tomancak, P.; Cardona, A. Fiji: An Open-Source Platform for Biological-Image Analysis. Nat Methods 2012, 9 (7), 676–682.

(25) Wickham, H. Ggplot2: Elegant Graphics for Data Analysis, 2nd ed.; Use R!; Springer International Publishing: Cham, Switzerland, 2016.

(26) Good, N. E.; Winget, G. D.; Winter, W.; Connolly, T. N.; Izawa, S.; Singh, R. M. Hydrogen Ion Buffers for Biological Research. Biochemistry 1966, 5 (2), 467–477.

(27) Lee, K. J.; Jordan, J. S.; Williams, E. R. Is Native Mass Spectrometry in Ammonium Acetate Really Native? Protein Stability Differences in Biochemically Relevant Salt Solutions. Anal Chem 2024, 96 (44), 17586–17593.

(28) LaCava, J.; Jiang, H.; Rout, M. P. Protein Complex Affinity Capture from Cryomilled Mammalian Cells. J Vis Exp 2016, No. 118. 10.3791/54518.

(29) Baldwin, R. L. How Hofmeister Ion Interactions Affect Protein Stability. Biophys J 1996, 71 (4), 2056–2063.

(30) Lo Nostro, P.; Ninham, B. W. Hofmeister Phenomena: An Update on Ion Specificity in Biology. Chem Rev 2012, 112 (4), 2286–2322.

(31) Gekko, K.; Timasheff, S. N. Mechanism of Protein Stabilization by Glycerol: Preferential Hydration in Glycerol-Water Mixtures. Biochemistry 1981, 20 (16), 4667–4676.

(32) Seddon, A. M.; Curnow, P.; Booth, P. J. Membrane Proteins, Lipids and Detergents: Not Just a Soap Opera. Biochim Biophys Acta 2004, 1666 (1-2), 105–117.

(33) Tjernberg, A.; Markova, N.; Griffiths, W. J.; Hallén, D. DMSO-Related Effects in Protein Characterization. J Biomol Screen 2006, 11 (2), 131–137.

(34) Yoshikawa, H.; Hirano, A.; Arakawa, T.; Shiraki, K. Effects of Alcohol on the Solubility and Structure of Native and Disulfide-Modified Bovine Serum Albumin. Int J Biol Macromol 2012, 50 (5), 1286–1291.

(35) Molecular Cloning, Expression, and Characterization of Amorpha-4,11-Diene Synthase, a Key Enzyme of Artemisinin Biosynthesis in Artemisia Annua L. Archives of Biochemistry and Biophysics 2000, 381 (2), 173–180.

(36) Bornhorst, J. A.; Falke, J. J. Purification of Proteins Using Polyhistidine Affinity Tags. Methods Enzymol 2000, 326, 245–254.

(37) Hochuli, E.; Döbeli, H.; Schacher, A. New Metal Chelate Adsorbent Selective for Proteins and Peptides Containing Neighbouring Histidine Residues. J Chromatogr 1987, 411, 177–184.

