## Supplementary information for "A miniaturized, high-throughput aqueous solvent-centric method for protein solubility screening"

### **Table S1.** Impact of the buffer composition on SDS-PAGE followed by coomassie blue staining from full E. coli bacterial lysates producing HISx8_Trx_Ruby2

| **Buffer Code** | **Buffer composition** | | | **SDS PAGE on bacterial lysate with HIS_Trx_Ruby2** |
| --- | --- | --- | --- | --- |
|  | **Buffer** | **Salt** | **Additive** |  |
| **A1** | Tris-HCl 50 mM; pH 8.5 | 50 mM NaCl | 10 % glycerol | +++ |
| **A3** | Tris-HCl 50 mM; pH 8.5 | 50 mM NaCl | 5 % ethanol | +++ |
| **A4** | Tris-HCl 50 mM; pH 8.5 | 50 mM NaCl | 1 % triton X10 | +++ |
| **A5** | Tris-HCl 50 mM; pH 8.5 | 50 mM NaCl | 100 mM Urea | +++ |
| **B2** | HEPES 50 mM; pH 7.0 | 50 mM NaCl | 5 % DMSO | +++ (inclined band) |
| **B3** | HEPES 50 mM; pH 7.0 | 50 mM NaCl | 5 % EtOH | +++ |
| **B4** | HEPES 50 mM; pH 7.0 | 50 mM NaCl | 1 % triton X10 | +++ (spread band) |
| **B5** | HEPES 50 mM; pH 7.0 | 50 mM NaCl | 100 mM Urea | +++ (inclined band) |
| **C** | Tris-HCl 50 mM; pH 8.5 | 300 mM NaCl | 10 % glycerol | + (inclined and blurred band) |
| **G4** | Tris-HCl 20 mM, pH 8.0 | 400 mM NaCl | 1 % triton X10 | - (blurred and spread band) |
| **H2** | Tris-HCl 20 mM, pH 8.0 | 0.4 M NH4Ac |  | - (blurred, unreadable band) |
| **H6** | Tris-HCl 20 mM, pH 8.0 | 2.0 M NH4Ac |  | - (blurred, unreadable band) |
| **J2** | Tris-HCl 20 mM, pH 8.0 | 0.4 M NH4Ac | 1 % triton X10 | + (spread and blurred band) |
| **K1** |  | 2.0 M NH4Ac |  | +++ |
| **K5** |  | 1.5 M NH4Ac |  | +++ (blurred band) |
| **P2** | HEPES 50 mM; pH 7.4 | 200 mM NaCl | 1 % triton X10 | + (spread band) |
| **R2** | HEPES 50 mM; pH 7.4 | 0.4 M NH4Ac |  | +++ (spread band) |
| **R6** | HEPES 50 mM; pH 7.4 | 2.0 M NH4Ac |  | +++ (blurred band) |
| **T2** | HEPES 50 mM; pH 7.4 | 0.4 M NH4Ac | 1 % triton X10 | - (spread, unreadable band) |
| **T6** | HEPES 50 mM; pH 7.4 | 2.0 M NH4Ac | 1 % triton X10 | +++ (bit lower intensity) |
| **W** | mQ water |  |  | +++ |

### **Table S2.** Impact of the buffer composition on SDS-PAGE followed by coomassie blue staining from full E. coli bacterial lysates HISx8_MBP_Ruby2

| **Buffer Code** | **Buffer composition** | | | **SDS PAGE on bacterial lysate with HIS_MBP_Ruby2** |
| --- | --- | --- | --- | --- |
|  | **Buffer** | **Salt** | **Additive** |  |
| **A1** | Tris-HCl 50 mM; pH 8.5 | 50 mM NaCl | 10 % glycerol | +++ |
| **A3** | Tris-HCl 50 mM; pH 8.5 | 50 mM NaCl | 5 % ethanol | +++ |
| **A4** | Tris-HCl 50 mM; pH 8.5 | 50 mM NaCl | 1 % triton X10 | +++ |
| **A5** | Tris-HCl 50 mM; pH 8.5 | 50 mM NaCl | 100 mM Urea | +++ |
| **B2** | HEPES 50 mM; pH 7.0 | 50 mM NaCl | 5 % DMSO | +++ |
| **B3** | HEPES 50 mM; pH 7.0 | 50 mM NaCl | 5 % EtOH | +++ (bit blurred band) |
| **B4** | HEPES 50 mM; pH 7.0 | 50 mM NaCl | 1 % triton X10 | +++ |
| **B5** | HEPES 50 mM; pH 7.0 | 50 mM NaCl | 100 mM Urea | +++ |
| **C** | Tris-HCl 50 mM; pH 8.5 | 300 mM NaCl | 10 % glycerol | +++ |
| **G4** | Tris-HCl 20 mM, pH 8.0 | 400 mM NaCl | 1 % triton X10 | - (blurred, unreadable band) |
| **H2** | Tris-HCl 20 mM, pH 8.0 | 0.4 M NH4Ac |  | - (blurred, unreadable band) |
| **H6** | Tris-HCl 20 mM, pH 8.0 | 2.0 M NH4Ac |  | - (blurred, spread, unreadable band) |
| **J2** | Tris-HCl 20 mM, pH 8.0 | 0.4 M NH4Ac | 1 % triton X10 | - (blurred, unreadable band) |
| **K1** |  | 2.0 M NH4Ac |  | - (blurred, unreadable band) |
| **K5** |  | 1.5 M NH4Ac |  | - (blurred, spread, unreadable band) |
| **P2** | HEPES 50 mM; pH 7.4 | 200 mM NaCl | 1 % triton X10 | + (blurred band) |
| **R2** | HEPES 50 mM; pH 7.4 | 0.4 M NH4Ac |  | +++ (bit blurred band) |
| **R6** | HEPES 50 mM; pH 7.4 | 2.0 M NH4Ac |  | + (spread, blurred band) |
| **T2** | HEPES 50 mM; pH 7.4 | 0.4 M NH4Ac | 1 % triton X10 | + (blurred band) |
| **T6** | HEPES 50 mM; pH 7.4 | 2.0 M NH4Ac | 1 % triton X10 | +++ (spread, blurred band) |
| **W** | mQ water |  |  | +++ |

### **Table S3.** Impact of the buffer composition on SDS-PAGE followed by coomassie blue staining from purified HISx8_Trx_Ruby2 protein.

| **Buffer Code** | **Buffer composition** | | | **SDS PAGE on purified HIS_Trx_Ruby2** |
| --- | --- | --- | --- | --- |
|  | **Buffer** | **Salt** | **Additive** |  |
| **A1** | Tris-HCl 50 mM; pH 8.5 | 50 mM NaCl | 10 % glycerol | +++ |
| **A3** | Tris-HCl 50 mM; pH 8.5 | 50 mM NaCl | 5 % ethanol | +++ |
| **A4** | Tris-HCl 50 mM; pH 8.5 | 50 mM NaCl | 1 % triton X10 | ++ (bit lower intensity) |
| **A5** | Tris-HCl 50 mM; pH 8.5 | 50 mM NaCl | 100 mM Urea | +++ |
| **B2** | HEPES 50 mM; pH 7.0 | 50 mM NaCl | 5 % DMSO | +++ |
| **B3** | HEPES 50 mM; pH 7.0 | 50 mM NaCl | 5 % EtOH | +++ |
| **B4** | HEPES 50 mM; pH 7.0 | 50 mM NaCl | 1 % triton X10 | +++ |
| **B5** | HEPES 50 mM; pH 7.0 | 50 mM NaCl | 100 mM Urea | +++ |
| **C** | Tris-HCl 50 mM; pH 8.5 | 300 mM NaCl | 10 % glycerol | +++ |
| **G4** | Tris-HCl 20 mM, pH 8.0 | 400 mM NaCl | 1 % triton X10 | +++ |
| **H2** | Tris-HCl 20 mM, pH 8.0 | 0.4 M NH4Ac |  | + (spread, blurred band) |
| **H6** | Tris-HCl 20 mM, pH 8.0 | 2.0 M NH4Ac |  | +++ (bit blurred band) |
| **J2** | Tris-HCl 20 mM, pH 8.0 | 0.4 M NH4Ac | 1 % triton X10 | - (blurred band) |
| **K1** |  | 2.0 M NH4Ac |  | +++ (bit blurred) |
| **K5** |  | 1.5 M NH4Ac |  | +++ |
| **P2** | HEPES 50 mM; pH 7.4 | 200 mM NaCl | 1 % triton X10 | +++ (bit blurred band) |
| **R2** | HEPES 50 mM; pH 7.4 | 0.4 M NH4Ac |  | + (spread blurred band) |
| **R6** | HEPES 50 mM; pH 7.4 | 2.0 M NH4Ac |  | +++ (blurred band) |
| **T2** | HEPES 50 mM; pH 7.4 | 0.4 M NH4Ac | 1 % triton X10 | +++ (bit blurred band) |
| **T6** | HEPES 50 mM; pH 7.4 | 2.0 M NH4Ac | 1 % triton X10 | +++ |
| **W** | mQ water |  |  | +++ |

### **Table S4.** Proteins expressed in this study

| Manuscript name | Database name | Expression vector | Experiment | Reference |
| --- | --- | --- | --- | --- |
| His8-Trx-mRuby2 | In-house | p3Xpress_Eco_Trx | Table 1 | This study |
| His8-MBP-mRuby2 | In-house | p3Xpress_Eco_MBP | Table 1, Figure 2 | This study |
| AMS | [Q9AR04](https://www.uniprot.org/uniprotkb/Q9AR04/entry), Uniprot | p3Xpress_Eco | Figure 3 | (21) |
| TPS1 | [A0A0E3NXY0](https://www.uniprot.org/uniparc/UPI000615CEEA/entry/A0A0E3NXY0), Uniprot | p3Xpress_Eco | Figure 3 | (21) |
| TPS2 | [A0A537EJD0](https://www.uniprot.org/uniparc/UPI001227849D/entry/A0A537EJD0), Uniprot | p3Xpress_Eco | Figure 3 | (21) |
| TPS3 | [A0A5E4I9B1](https://www.uniprot.org/uniparc/UPI0011C31E89/entry/A0A5E4I9B1), Uniprot | p3Xpress_Eco | Figure 3 | (21) |
| TPS4 | [A0A2H0W7N1](https://www.uniprot.org/uniprotkb/A0A2H0W7N1/entry), Uniprot | p3Xpress_Eco | Figure 3 | (21) |
| TPS5 | [A0A450SFA8](https://www.uniprot.org/uniprotkb/A0A450SFA8/entry), Uniprot | p3Xpress_Eco | Figure 3 | (21) |
| TPS6 | [A0A5S9IQ85](https://www.uniprot.org/uniprotkb/A0A5S9IQ85/entry), Uniprot | p3Xpress_Eco | Figure 3 | (21) |
| TPS7 | UPI00031437D0 | p3Xpress_Eco | Figure 3 | (21) |
| TPS8 | [M2QMG2](https://www.uniprot.org/uniprotkb/M2QMG2/entry), Uniprot | p3Xpress_Eco | Figure 3 | (21) |
| Protein1 | In-house | pET30a(+) | Figure 4 | This study |
| Protein2 | In-house | pET30a(+) | Figure 4 | This study |
| Protein3 | In-house | pET30a(+) | Figure 4 | This study |
| Protein4 | In-house | pET30a(+) | Figure 4 | This study |
| Protein5 | In-house | pET30a(+) | Figure 4 | This study |

*De novo* generated proteins **1-5.**

**1:**

MQTILQDYANVVDGSFFRHWKLFSGMEAKNLQRLYNGLVERECEDGQEYSLLDQRSKAHIDSIALSIYKHVTSHEFSLPSWVGLASDMLTGGTLRVVAGGLQVKTINNPH

**2:**

MEKFLPNYLKDVGGSLSRYREFICEIVCENQQRHYKRSFELQCKEILHDILLEQVTKPDVYNRDLAIDELATSHAFSMLSLPILETEIQIGGIINMITERVDLTKVNDPRG

**3:**

MQTFLQAYSTVGEVCLFRYRQWFFAMQDKKLMKDYKRLFEVECKETQHDGLLDRATKPNRDNVDVALAKLASTDALSLAPWLGLDSGILIADIMHMRTEGLHPNTRYSPPG

**4:**

MQMILQKYTDVVEVRSFNYWQRFTEMVSEKQKKYAKGFAAWECEKALEDTLLDTLAKPQFSKVDAGSRKLSNARELSLLPLPIVATAIQVAATSKAITEGLHLTKVYDLRG

**5**:

MEKFFQTNTVVVDVRSYKYWERFSAMVDKNLARVYKGPDEWEYKDSQEDGMLEIRPKANMANVDLSTGELAKILVLRLPLSASVDFGMQIGDIMRLMSDRLEADKINDLHG

### **Figure S1.** SDS-PAGE with coomassie blue staining from E. coli *bacterial lysates producing HISx8_Trx_Ruby2.* W is extraction with water. Letters correspond to the buffer code.

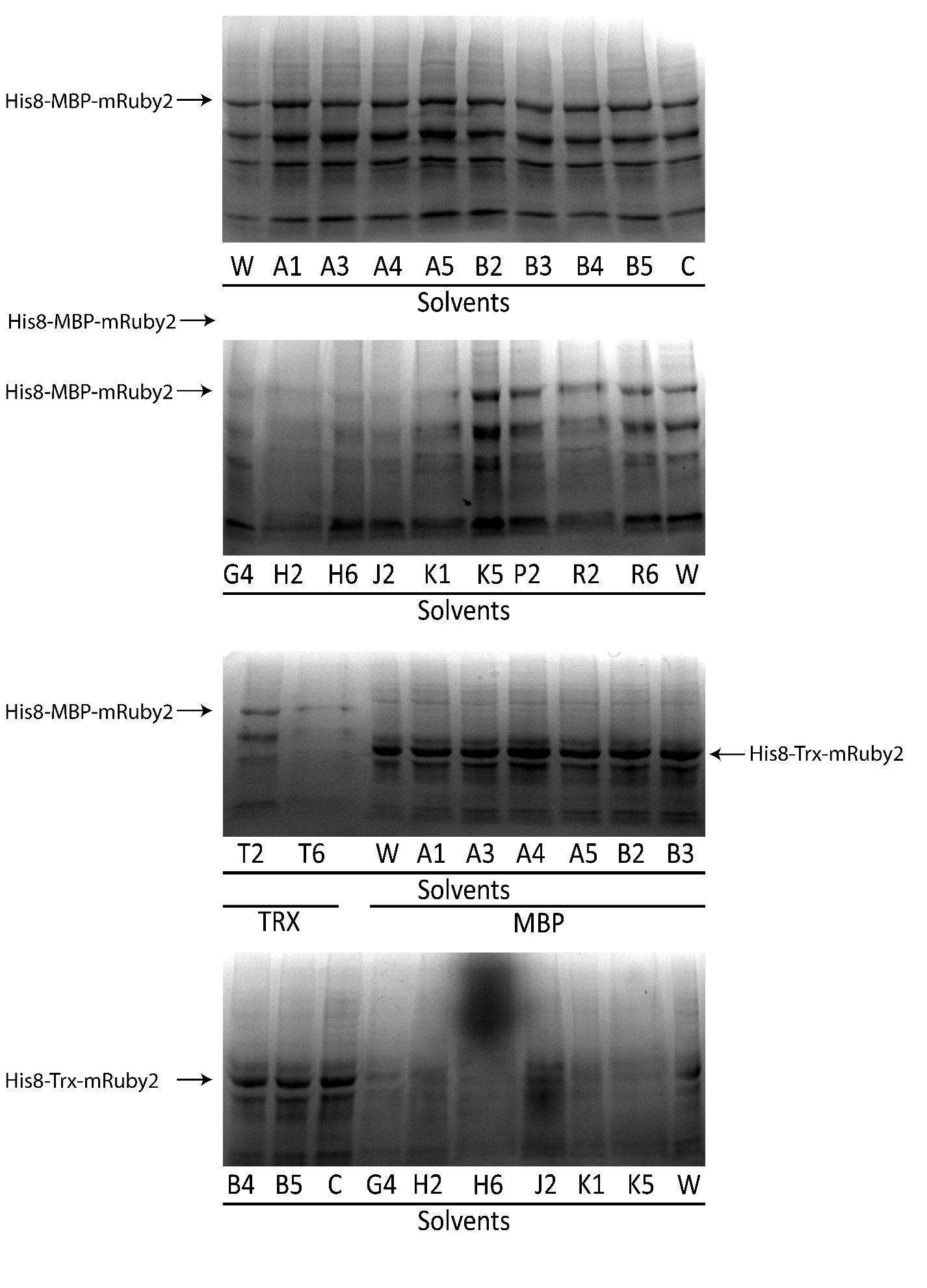

### **Figure S2.** SDS-PAGE with coomassie blue staining from E. coli *bacterial lysates producing HISx8_Trx_Ruby2.* W is extraction with water. Letters correspond to the buffer code.

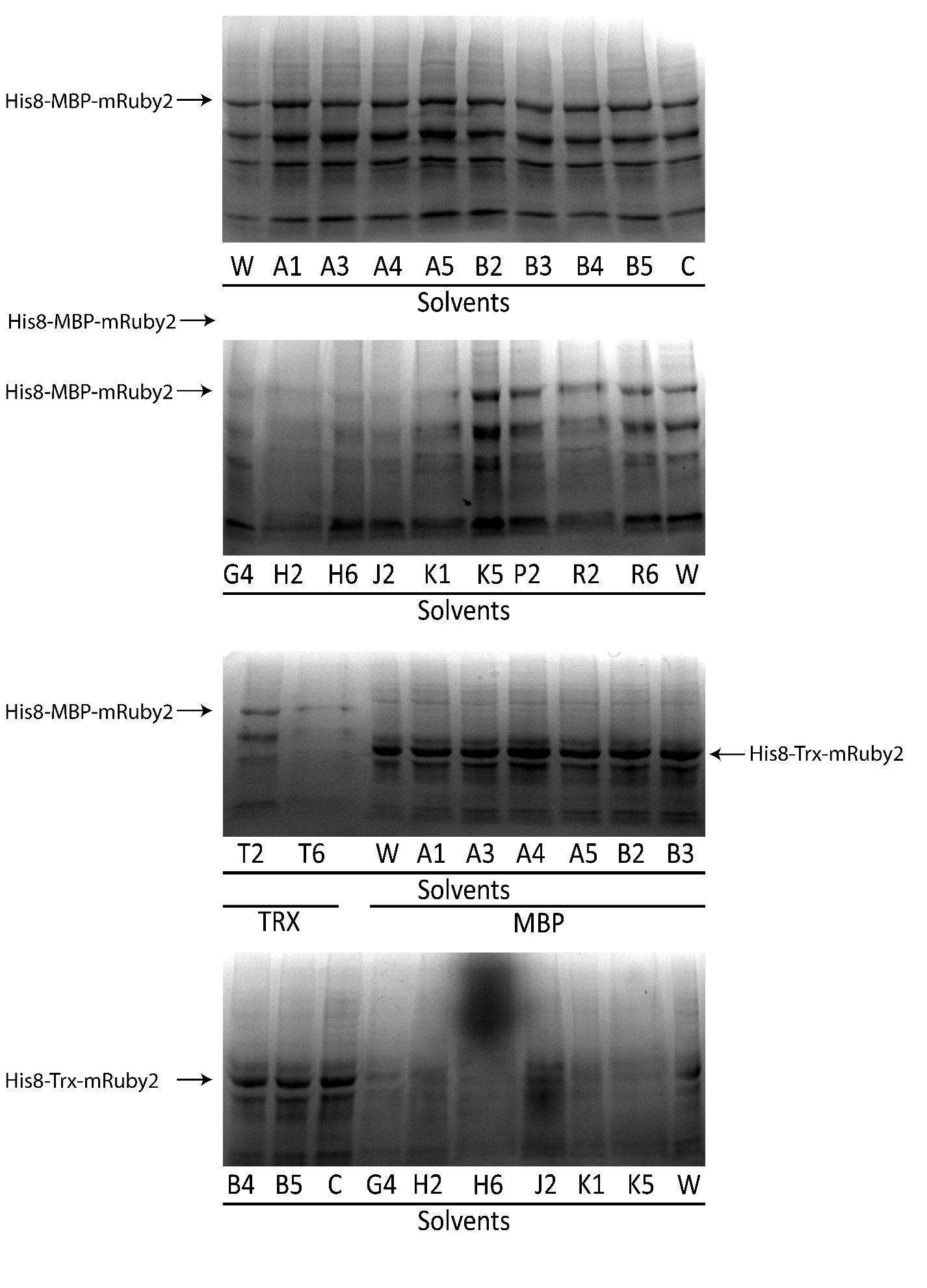

### **Figure S3**. SDS-PAGE with coomassie blue straining from E. coli *bacterial lysates producing HISx8_Trx/MBP_Ruby2.* W is extraction with water. Letters correspond to the buffer code.

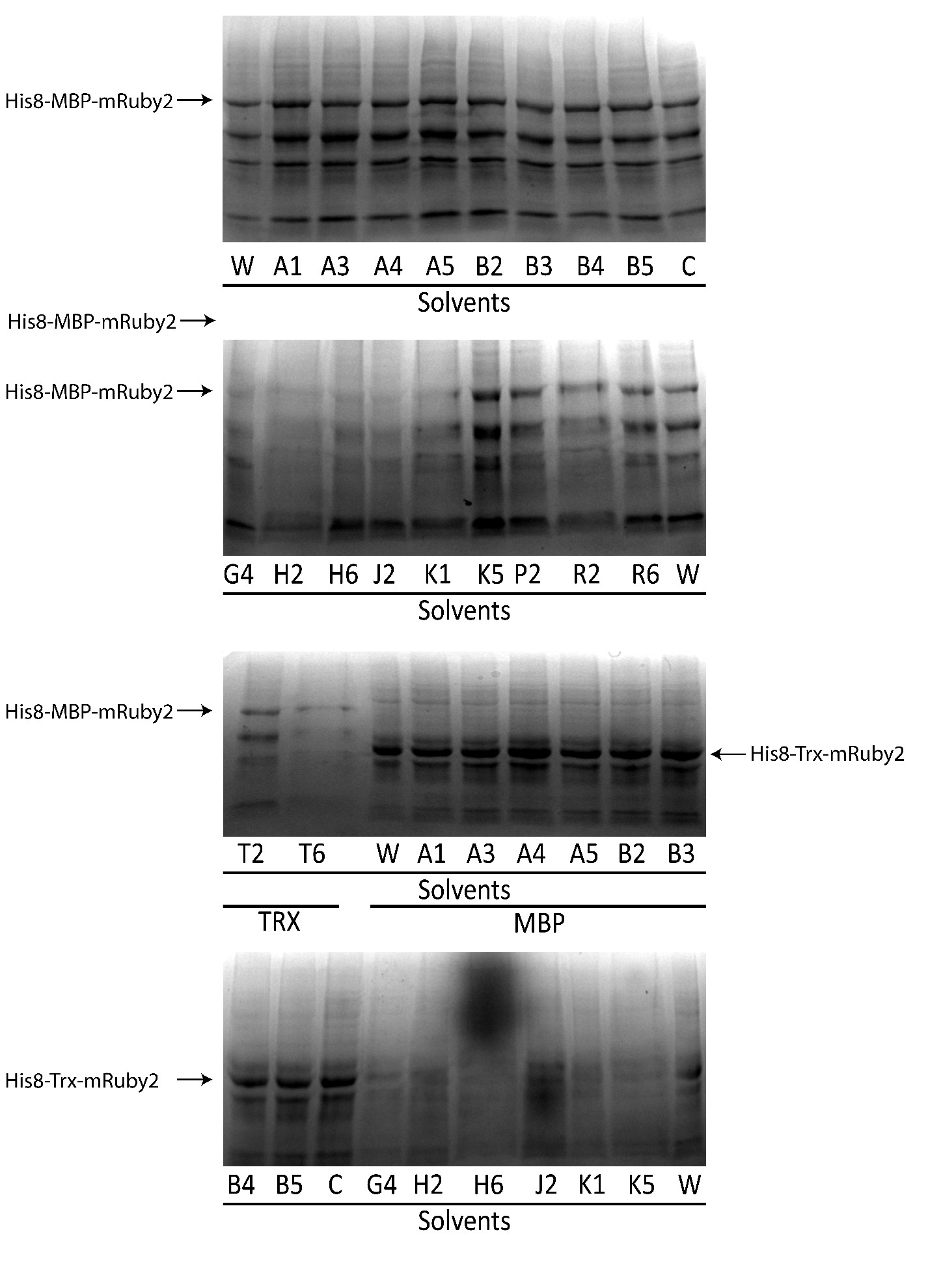

### **Figure S4**. SDS-PAGE with coomassie blue staining from E. coli *bacterial lysates producing HISx8_MBP_Ruby2.* W is extraction with water. Letters correspond to the buffer code.

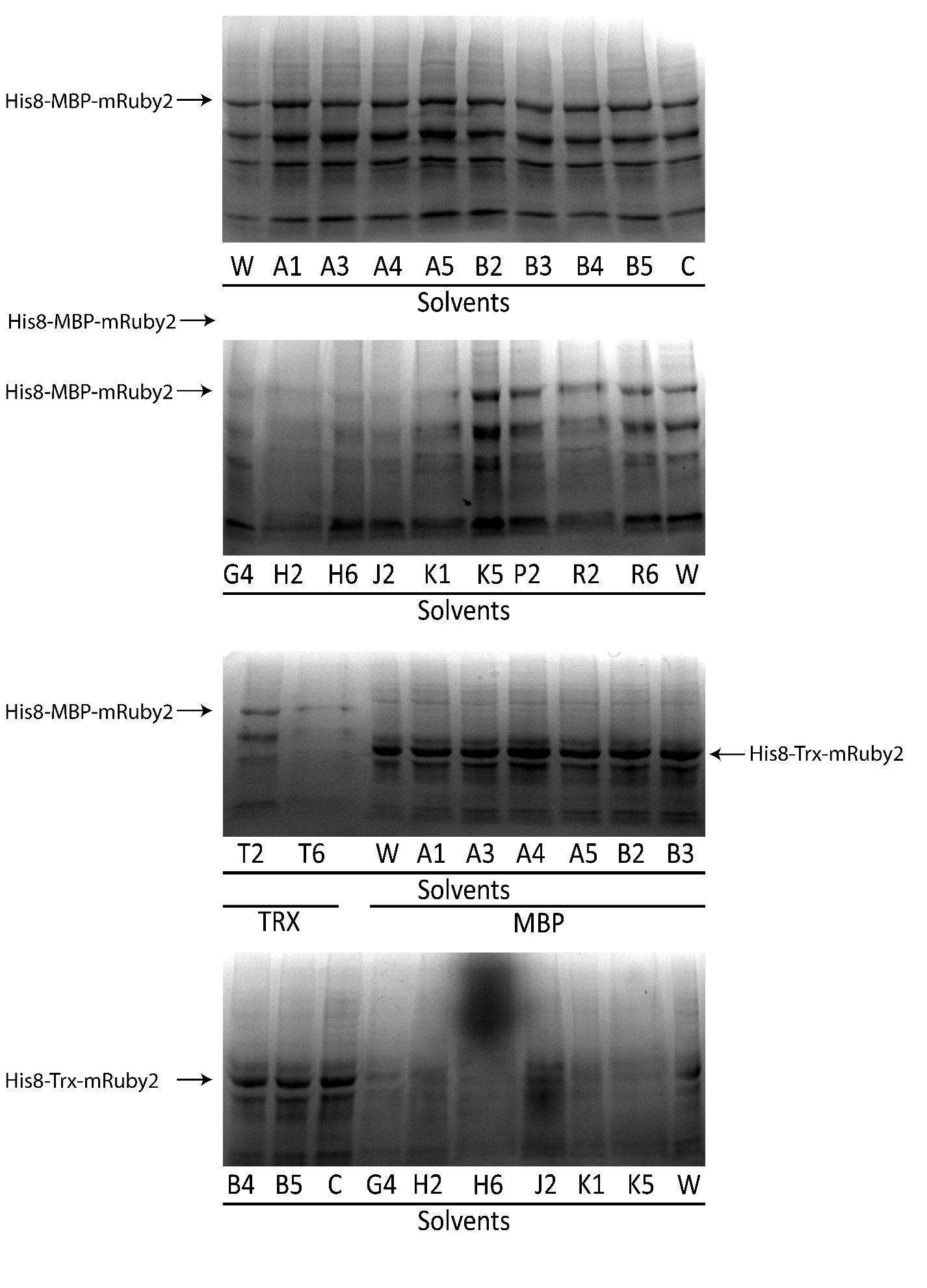

### **Figure S5**. SDS-PAGE with coomassie blue staining from E. coli *bacterial lysates producing HISx8_MBP_Ruby2.* W is extraction with water. Letters correspond to the buffer code.

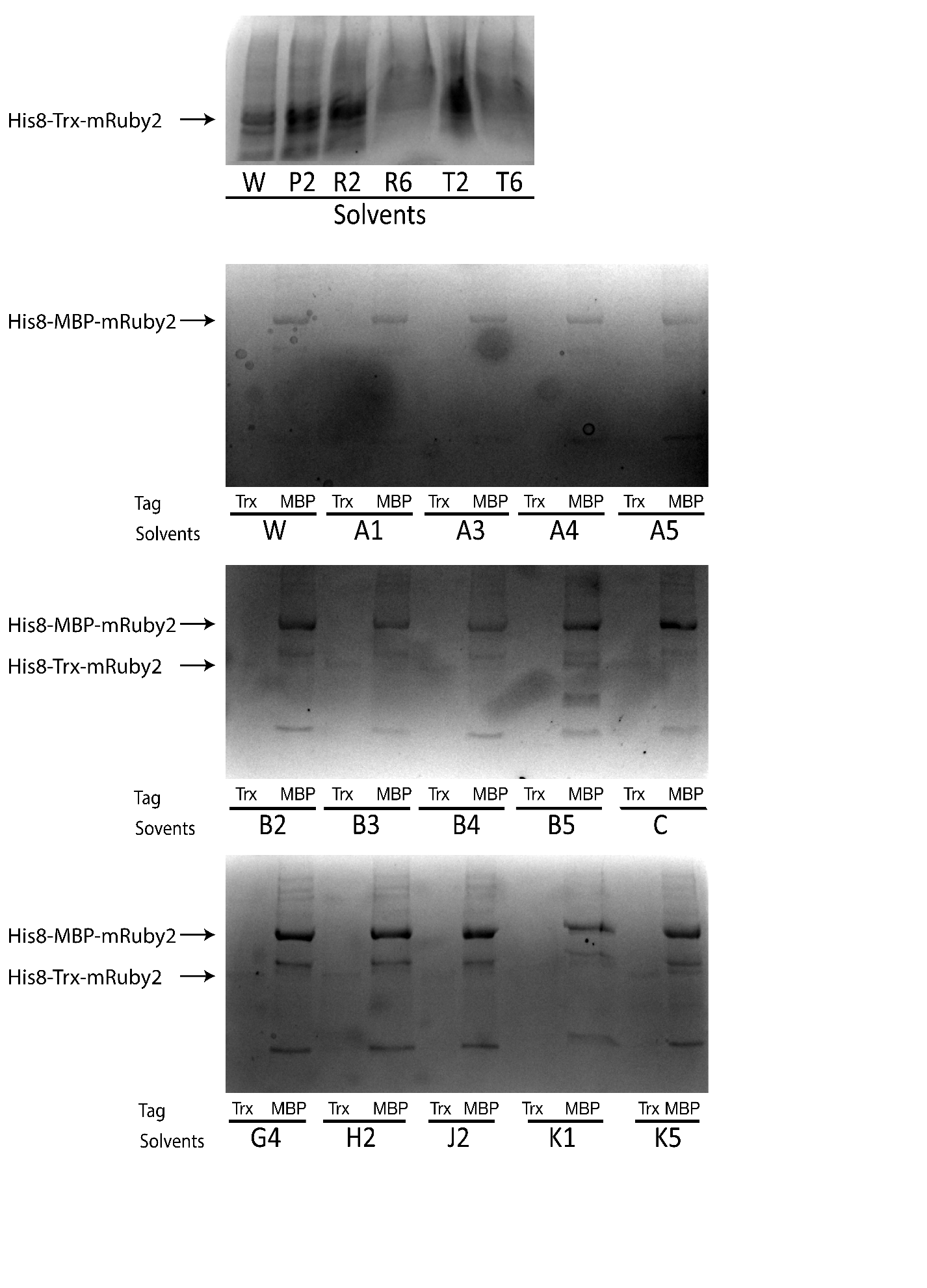

### **Figure S6**. SDS-PAGE with coomassie blue staining from purified *HISx8_Trx/MBP_Ruby2 protein.* W is extraction with water. Letters correspond to the buffer code.

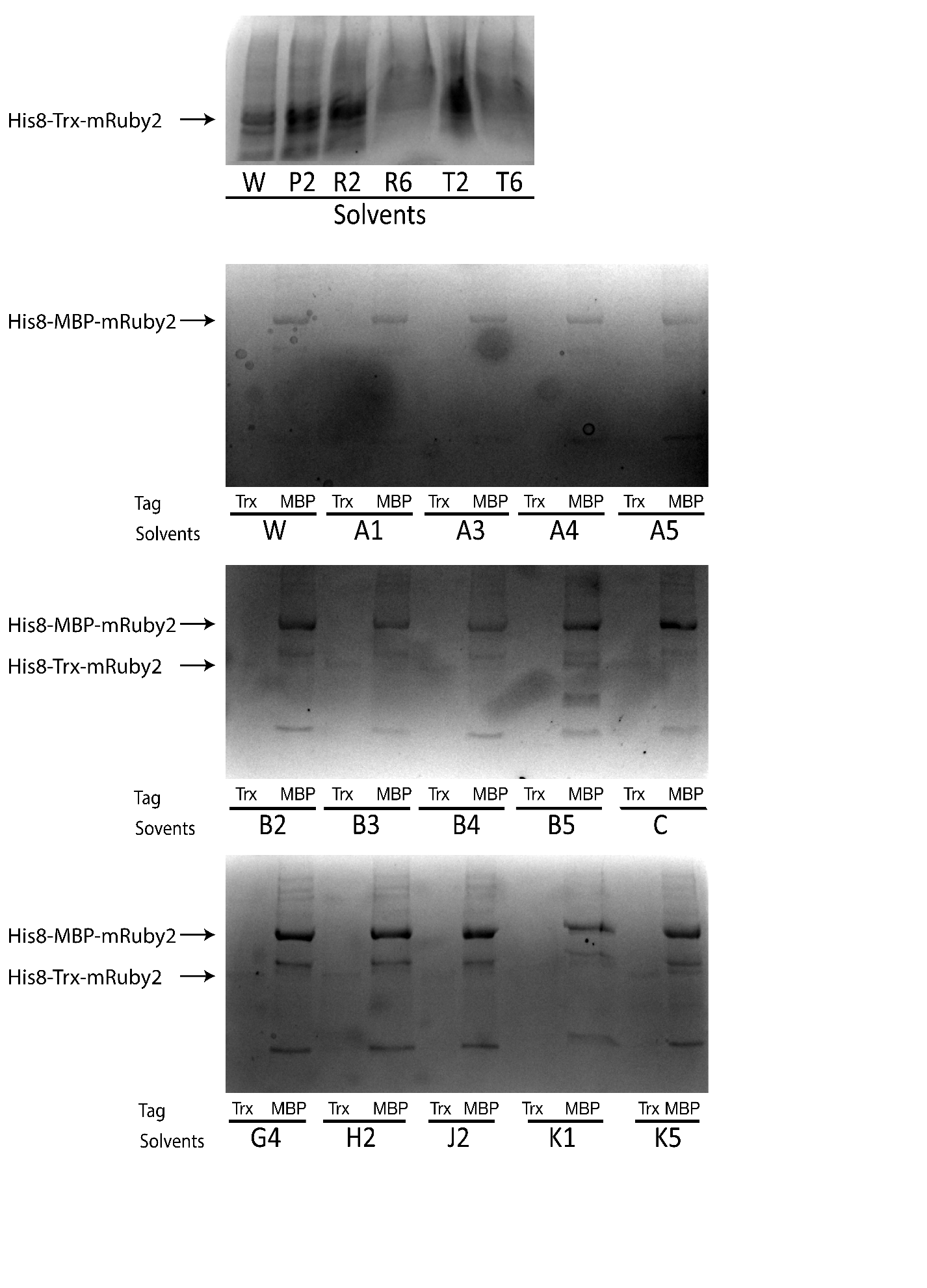

### **Figure S7**. SDS-PAGE with coomassie blue staining from purified *HISx8_Trx/MBP_Ruby2 protein.* W is extraction with water. Letters correspond to the buffer code.

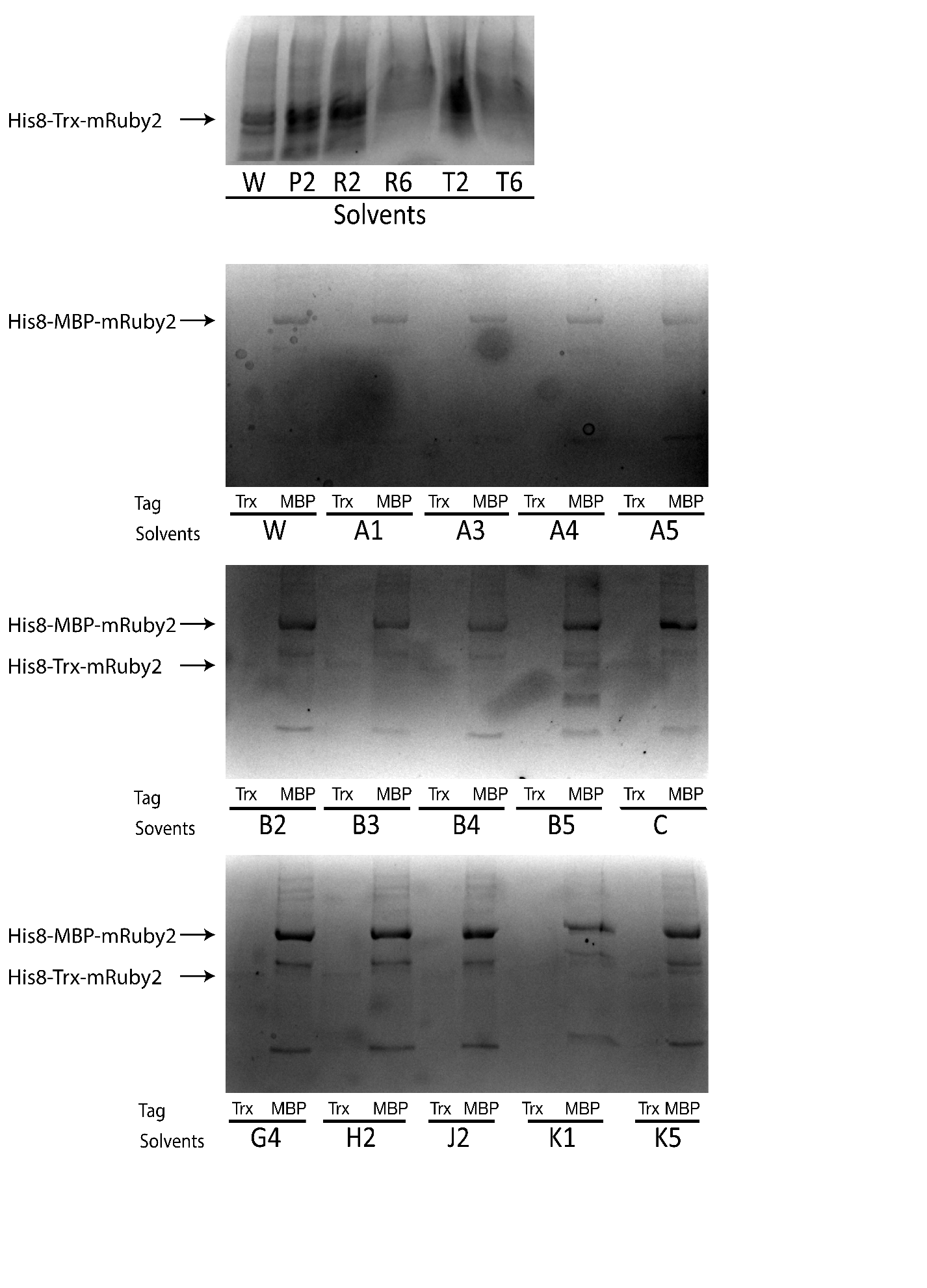

### **Figure S8**. SDS-PAGE with coomassie blue staining from purified *HISx8_Trx/MBP_Ruby2 protein.* W is extraction with water. Letters correspond to the buffer code.

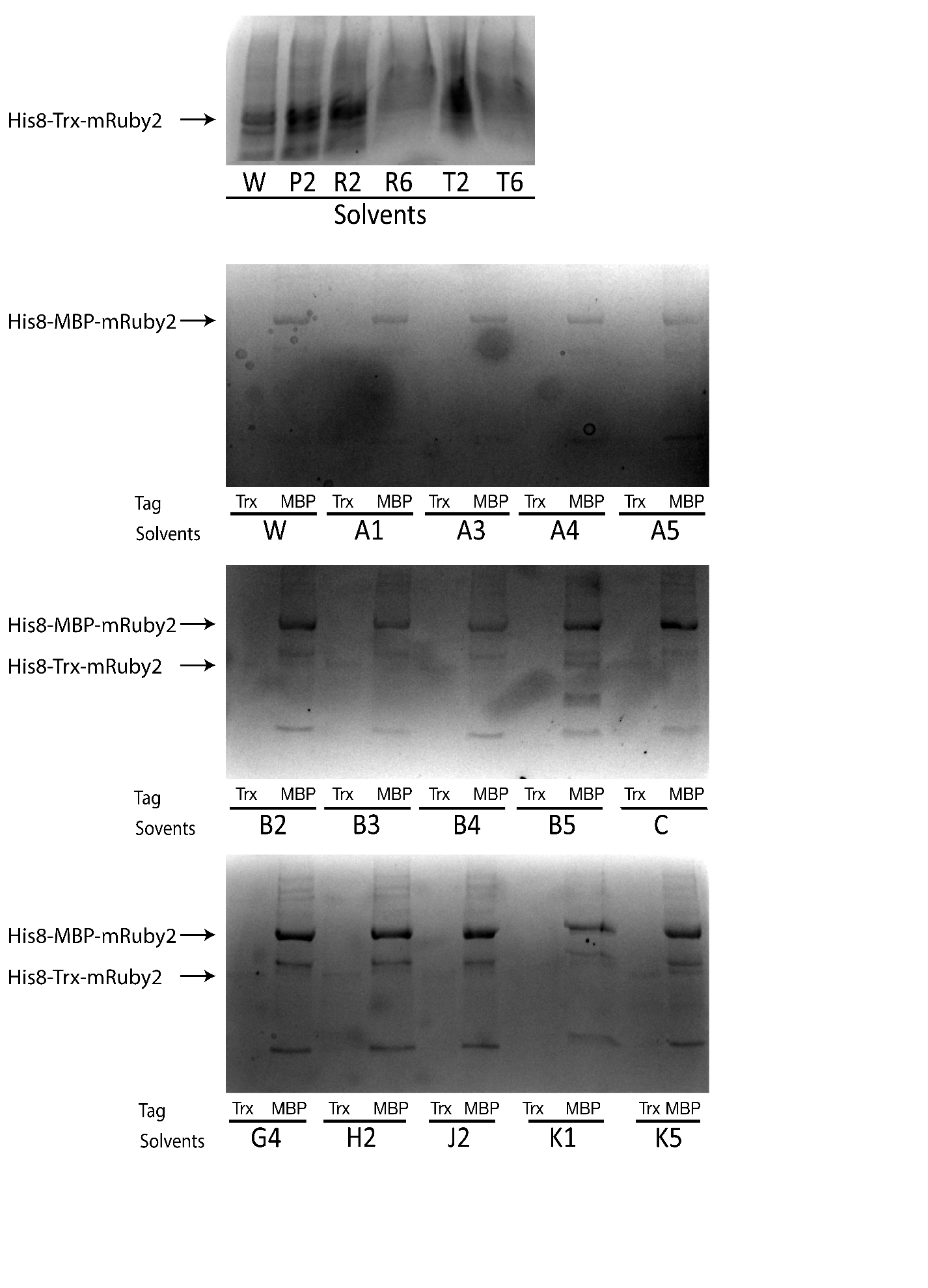

### **Figure S9**. SDS-PAGE with coomassie blue staining from purified *HISx8_Trx/MBP_Ruby2 protein.* W is extraction with water. Letters correspond to the buffer code

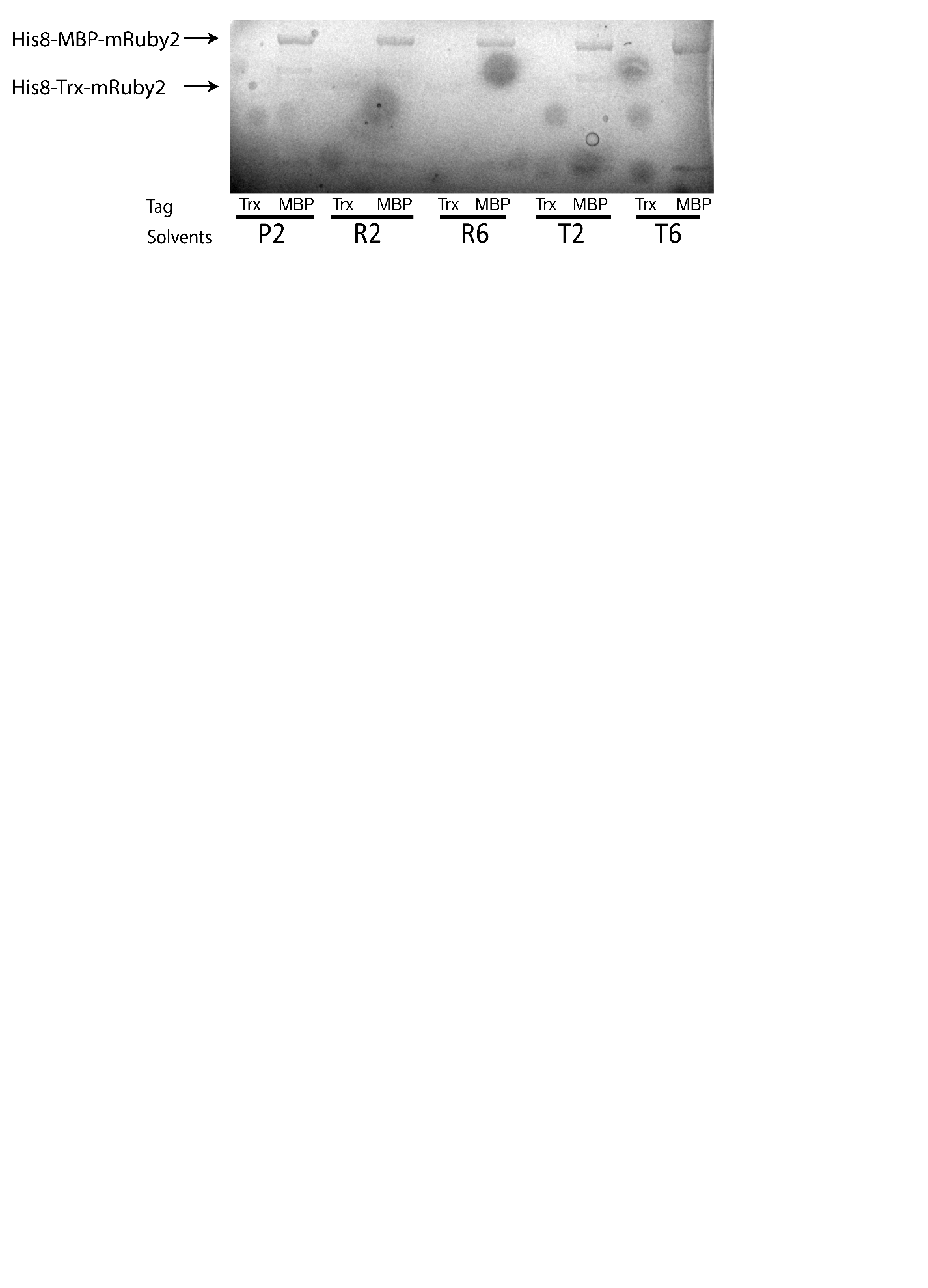

### **Figure S10**. Fluorescence-lifetime imaging microscopy for protein 1, 2 and 5

**Protein 1**.

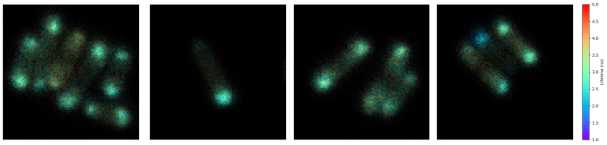

**Protein 2.**

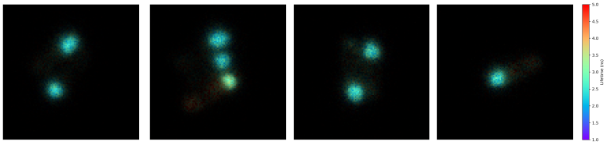

**Protein 5.**

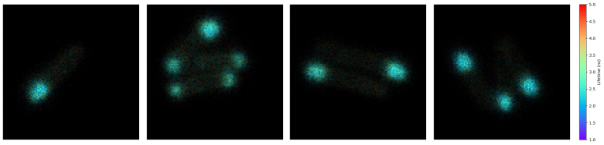

### **Figure S11**. Extraction of protein 3 by chemical lysis. T: total extract, S: soluble fraction.

Protein 3.

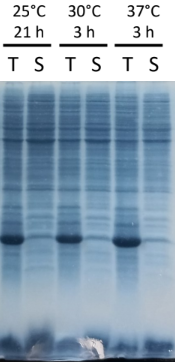
